# Cortical and thalamic connectivity of occipital visual cortical areas 17, 18, 19 and 21 of the domestic ferret (Mustela putorius furo)

**DOI:** 10.1101/491399

**Authors:** Leigh-Anne Dell, Giorgio M. Innocenti, Claus C. Hilgetag, Paul R. Manger

## Abstract

The present study describes the ipsilateral and contralateral cortico-cortical and cortico-thalamic connectivity of the occipital visual areas 17,18, 19 and 21 in the ferret using standard anatomical tract-tracing methods. In line with previous studies of mammalian visual cortex connectivity, substantially more anterograde and retrograde label was present in the hemisphere ipsilateral to the injection site compared to the contralateral hemisphere. Ipsilateral reciprocal connectivity was the strongest within the occipital visual areas, while weaker connectivity strength was observed in the temporal, suprasylvian and parietal visual areas. Callosal connectivity tended to be strongest in the homotopic cortical areas, and revealed a similar areal distribution to that observed in the ipsilateral hemisphere, although often less widespread across cortical areas. Ipsilateral reciprocal connectivity was observed throughout the visual nuclei of the dorsal thalamus, with no contralateral connections to the visual thalamus being observed. The current study, along with previous studies of connectivity in the cat, identified the posteromedial lateral suprasylvian visual area (PMLS) as a distinct network hub external to the occipital visual areas in carnivores, implicating PMLS as a potential gateway to the parietal cortex for dorsal stream processing. These data will also contribute to the Ferretome (www.ferretome.org), a macro connectome database of the ferret brain, providing essential data for connectomics analyses and cross-species analyses of connectomes and brain connectivity matrices, as well as providing data relevant to additional studies of cortical connectivity across mammals and the evolution of cortical connectivity variation.

## Introduction

It is well established that visual information is processed in the primary and associative visual cortical areas of mammals, and that these areas are embedded in an extensive network of cortical and subcortical connections (e.g. van Essen, Andersen, & Felleman, 1992; Ungerleider & Haxby, 1994; Scannell, Blakemore, & Young, 1995). The organization of such networks provides a structural basis for understanding of how visual information is perceived and used to initiate behaviour (e.g. Goodale & Milner, 1992). Anatomical tract-tracing, in conjunction with electrophysiological mapping studies, has been the primary tool used to provide insights into the visual cortical network organization (e.g. Symonds & Rosenquist, 1984; Young, Scannell, Burns, & Blakemore, 1994; Vanduffel, Payne, Lomber, & Orban, 1997; de Pasquale & Sherman, 2011; Markov et al., 2014). While diffusion tensor imaging (DTI) has gained popularity in recent years, this technique is complicated by numerous measurement and artifact problems and cannot, currently, provide the structural specificity obtained by anatomical tract-tracing methods (Vanduffel et al., 1997; van Essen et al., 2012; Meier-Hain et al., 2017). Anatomical tract-tracing has been used to examine connectivity of the visual cortex in a variety of species (e.g. mouse, Wang & Burkhalter, 2007; rat, Coogan & Burkhalter, 1990; cat, Symonds & Rosenquist, 1984; squirrel monkey, Tigges J, Tigges M, Anschel, Cross, Letbetter, & McBride, 1981; macaque monkey, Rockland & Pandya, 1979, Callaway 1998, Stettler, Das, Bennett, & Gilbert, 2002); however, at present such studies in the ferret are limited. The global callosal connectivity of the ferret visual cortex, and its relation to retinotopy within the various visual cortical areas, has been examined (Innocenti, Manger, Masiello, Colin, & Tettoni, 2002; Manger, Kiper, Masiello, Murillo, Tettoni, Hunyadi, & Innocenti, 2002a; Manger, Masiello, & Innocenti, 2002b; Manger, Nakamura, Valentiniene, & Innocenti, 2004), as well as the connectivity of the cortex with the superior colliculus (Manger, Restrepo, & Innocenti, 2010) and claustrum (Patzke, Innocenti, & Manger, 2014), but to date only one study has examined the cortico- cortical connectivity of a specific visual cortical area, the posteromedial lateral suprasylvian area (PMLS), in the ferret (Cantone, Xiao, & Levitt, 2006).

Over the past three decades the ferret has become an increasingly used animal model for studying the visual system, but while the location and properties of several visual cortical areas have been described (Innocenti et al., 2002, Manger et al., 2002a,b, 2004; Manger, Engler, Moll, & Engel 2005, 2008; Cantone et al., 2006; Homman-Ludiye, Manger, & Bourne, 2010), we do not fully understand information flow between and within different areas of the visual cortex or how these different cortical areas are connected with each other and with subcortical structures in the ferret. This information is pivotal to using the ferret as an animal model in vision research, and also for cross-species analyses of connectomes and brain connectivity matrices. Thus, this paper serves to initiate our systematic analysis of the connectivity of the ferret visual cortex by examining cortico-cortical and cortico-thalamic connectivity of the occipital visual areas 17, 18, 19 and 21 using standard anatomical tract-tracing methods. This data will also contribute to the Ferretome (www.ferretome.org), a macroconnectome database of the ferret brain (Sukhinin, Engel, Manger, & Hilgetag, 2016), to facilitate initiatives aimed at understanding how the brain processes and perceives visual information that can be used to guide and direct behaviour.

## Material and Methods

### Surgical procedure and tracer injections

Eight adult female ferrets (*Mustela putorius*), weighing between 600g and 1000g, were used in the current study (two injection sites per cortical area). The experiments were conducted according to the Swedish and European Community guidelines for the care and use of animals in scientific experiments. All animals were initially anesthetized with i.m. doses of ketamine hydrochloride (Ketalar, 10mg/kg) and medetomidin hydrochloride (Domitor, 0.08mg/kg), supplemented with atropine sulphate (0.15mg/kg) and placed in a stereotaxic frame. A mixture of 1% isoflurane in a 1:1 nitrous oxide and oxygen mixture was delivered through a mask while the animal maintained its own respiration. Anesthetic level was monitored using the eye blink and withdrawal reflexes, in combination with heart rate measurement. The occipital cortex was exposed under aseptic conditions and in each animal numerous (fewer than 20) electrophysiological recordings were taken to ensure correct placement of the tracer within a specific cortical area near the representation of the horizontal meridian (Manger et al., 2002a,b, 2004, 2005). Approximately 500 nl of tracer (biotinylated dextran amine, BDA 10 k, 5% in 0.1 M phosphate buffer; Molecular Probes) was delivered at each injection site using a Hamilton microsyringe (Figs. 1, 2a, 2b). After the completion of the injections, a soft contact lens was cut to fit over the exposed cortex, while the retracted dura mater was pulled over the contact lens and the excised portion of bone repositioned and held in place with dental acrylic. The temporal muscle was reattached using surgical glue and the midline incision of the skin sutured. Antibiotics were administered to all cases (Terramycin, 40 mg/kg, daily for 5 days) and these animals were given a 2-week recovery period to allow for tracer transport. At the end of this period, the animals were euthanized with a lethal dose of sodium pentobarbital (80 mg/kg, i.p.) and perfused intracardially, initially with a rinse of 0.9% saline (4°C, 500ml/kg), followed by fixation with 4% paraformaldehyde in 0.1M phosphate buffer (4°C, 1000 ml/kg).

**Figure 1:**
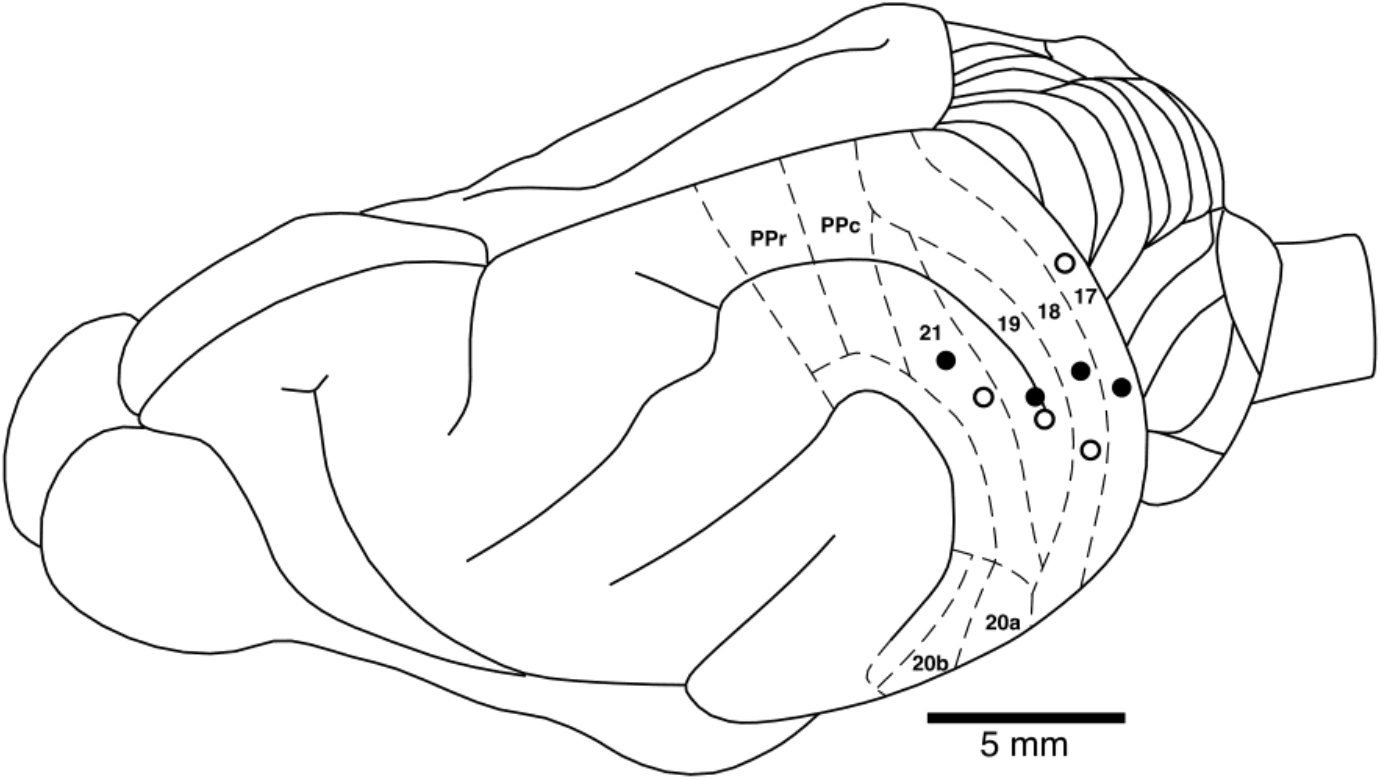
Locations of the occipital visual areas injection sites analyzed in the current study in relation to many of the known boundaries of visual cortical areas in a dorsolateral view of the ferret brain. Closed circles represent the injections sites where the brain was sectioned in a coronal plane. Open circles represent the injections sites where the cerebral cortex was manually semi-flattened for analysis. See list for abbreviations.

**Figure 2:**
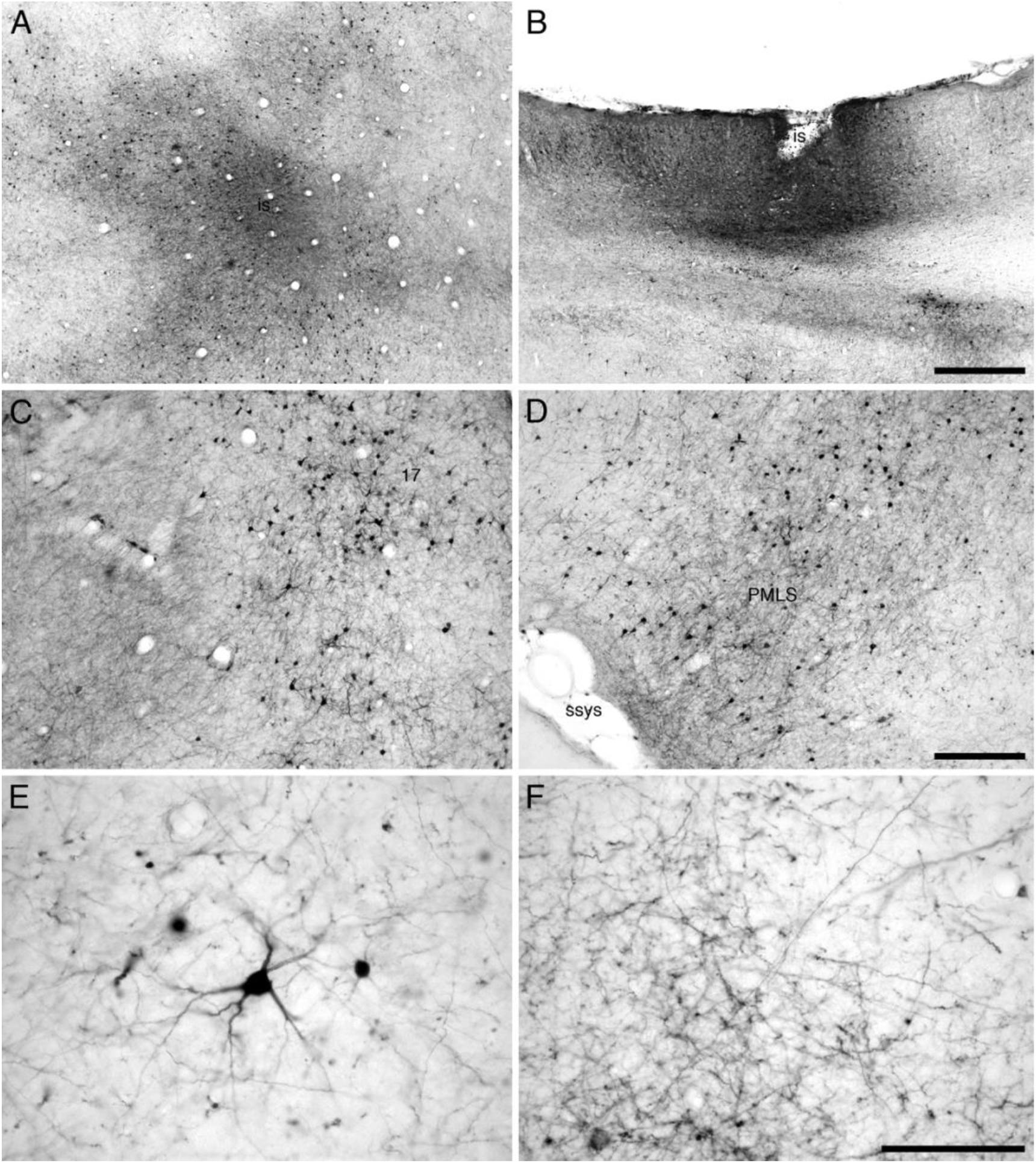
Photomicrographs showing examples of injection sites (**a, b**) and labelled cells (**c-f**) that were analyzed in the current study. (**a**) Injection site (**is**) in area **19** of a semi-flattened cerebral cortex showing the spread of tracer around the injection site, approximately a 500 μm halo of tracer, indicating the areal specificity of the injections made in the current study. (**b**) Injection site (**is**) in area **18** from a coronal section through the cerebral cortex, showing the spread of tracer and the limitation of the injection site to the cerebral cortex and not extending into the underlying white matter. Labelled cells and terminals in area **17** (**c**) and the posteromedial lateral suprasylvian area (**PMLS, d**) following transport from the injection site depicted in a. (e) High magnification image of a retrogradely labelled cell in area **21** following transport from the injection site depicted in **a**. (**f**) High magnification image of anterogradely labelled axons in the posterior parietal cortex (area **PPc**) following transport from the injection site depicted in **a**. Scale bar in **b** = 500 μm and applies to **a** and **b**. Scale bar in **d** = 250 μm and applies to **c** and **d**. Scale bar in **f** = 100 μm and applies to **e** and **f**. **ssys** – suprasylvian sulcus. In images **a, c-f**, the midline of the brain is to the top of the image and rostral to the left. In image **b**, the pial surface has been rotated to the top of the image, with the midline of the brain to the right.

### Sectioning and staining procedures

The brains were removed from the skull and post-fixed overnight in 4% paraformaldehyde in 0.1M phosphate buffer and then transferred to a 30% sucrose solution in 0.1M phosphate buffer (4°C) and allowed to equilibrate. The brains were either: (1) frozen in dry ice and sectioned at 50 μm on a freezing microtome in a coronal plane (4 cases, one for each of the occipital cortical areas) for a one in four series for Nissl (cresyl violet), myelin (Gallyas, 1979), cytochrome oxidase (Carroll & Wong-Riley, 1984) and BDA; or (2) cryoprotected and the cerebral cortex dissected away from the remainder of the brain and the dorsolateral surface semi-flattened (4 cases, one for each of the occipital cortical areas) between two glass slides, frozen onto the cold microtome stage and sectioned parallel to the semi-flattened surface at 50 μm for a one in two series for BDA and cytochrome oxidase. For BDA tracer visualization, the sections were incubated in 0.5% bovine serum albumin in 0.05M Tris buffer for 1h, followed by incubation in an avidin-HRP solution for 3h. A 10 min pre-incubation in 0.2% NiNH_4_SO_4_ preceded the addition of H_2_O_2_ (200 μl/l) to this solution, at which time the sections were monitored visually for the reaction product. To stop the reaction the sections were placed in 0.05 M Tris buffer. All sections were mounted on 0.5% gelatine coated slides, dehydrated in a graded series of alcohols, cleared in xylene and coverslipped with Depex mounting medium. All injection sites resulted in robust anterograde and retrograde transport of tracer in both the cerebral cortex (Fig. 2c-f) and the visual portion of the dorsal thalamus (Fig. 3).

**Figure 3:**
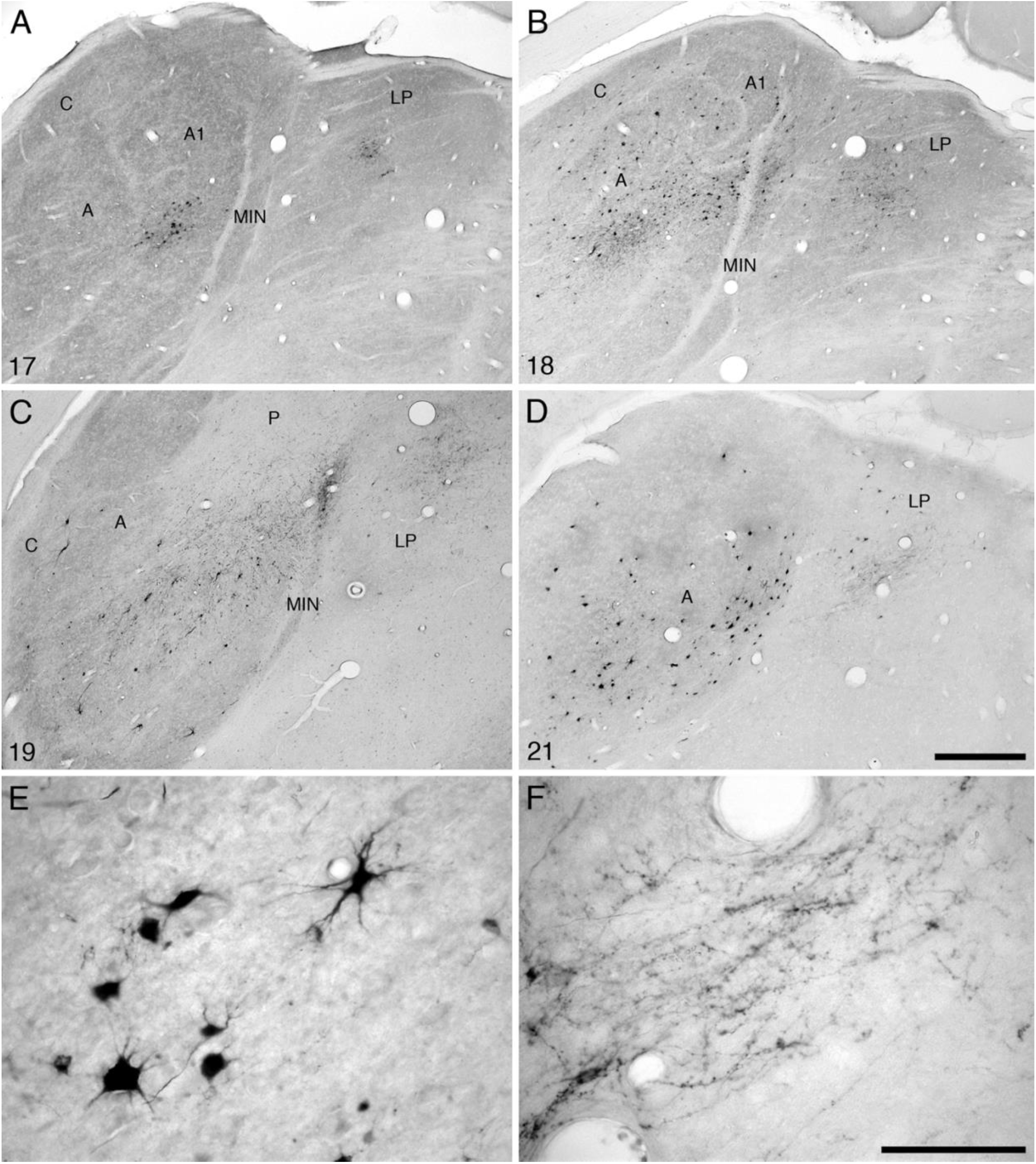
Photomicrographs showing examples of retrogradely labelled cells and anterogradely labelled axons in the visual thalamus of the ferret following injections into the occipital visual areas **17** (**a**), **18** (**b**), **19** (**c**) and **21** (**d**). (**e**) High magnification image of retrogradely labelled cells in the lateral geniculate nucleus following transport from the injection site in area **21**. (**f**) High magnification image of anterogradely labelled axons in the lateral posterior nucleus following transport from the injection site in area **21**. Scale bar in **d** = 500 μm and applies to **a – d**. Scale bar in **f** = 100 μm and applies to **e** and **f**. In all images dorsal is to the top and medial to the right. See list for abbreviations.

### Qualitative and quantitative analysis

For qualitative analysis, the stained sections were examined under low and high power magnification using a light microscope to determine in which sections through the cortex labelled cell bodies and terminals were present. Under low power stereomicroscopy using the flattened sections, the edges of each section were drawn with the aid of a camera lucida, and the location of the injection site marked. Areal borders were delineated and drawn using the cytochrome oxidase stained sections. The sections reacted for BDA were then matched to these drawings and the locations of the individual retrogradely labelled cells plotted and regions of anterogradely labelled axonal terminals demarcated. The drawings were scanned and redrawn using a Canvas X Pro 16 drawing program (ACD Systems International Inc., USA). Digital photomicrographs were captured using a Zeiss Axioskop and the Axiovision software. No pixilation adjustments or manipulation of the captured images were undertaken, except for contrast and brightness adjustment using Adobe Photoshop 7.

To quantify the retrograde BDA labelling per injection site, and control for variance in the size of the injection, a fraction of labelled neurons (N%, Table 1) was calculated using the formula:

**Table 1:** Fraction of retrogradely labelled neurons (N%), in the various visual cortical areas of the ipsilateral and contralateral hemispheres of the ferret following injections of biotinylated dextran amine (BDA) into the occipital visual areas **17, 18**, **19** and **21**. See list for abbreviations.

|  | Occipital |  |  |  | Temporal |  |  | Suprasylvian |  |  |  |  |  | Parietal |  |
| --- | --- | --- | --- | --- | --- | --- | --- | --- | --- | --- | --- | --- | --- | --- | --- |
| Injection site | 17 | 18 | 19 | 21 | 20a | 20b | PS | PMLS | PLLS | AMLS | ALLS | VLS | DLS | PPc | PPr |
| Ipsilateral connectivity |  |  |  |  |  |  |  |  |  |  |  |  |  |  |  |
| 17 | 38.25 | 32.60 | 15.36 | 5.01 | 0.00 | 0.00 | 0.00 | 3.13 | 0.00 | 0.00 | 0.00 | 0.00 | 0.00 | 0.00 | 0.00 |
| 18 | 7.80 | 25.94 | 22.18 | 21.59 | 2.99 | 3.96 | 0.30 | 3.80 | 0.42 | 0.08 | 0.00 | 1.94 | 0.00 | 0.00 | 0.00 |
| 19 | 8.56 | 19.38 | 24.32 | 18.06 | 2.07 | 1.40 | 0.19 | 7.92 | 0.00 | 0.81 | 0.00 | 1.23 | 0.00 | 1.34 | 0.08 |
| 21 | 7.13 | 16.77 | 24.45 | 27.25 | 4.92 | 3.49 | 0.66 | 6.31 | 0.49 | 0.82 | 0.11 | 1.50 | 0.02 | 2.65 | 0.20 |
| Contralateral connectivity |  |  |  |  |  |  |  |  |  |  |  |  |  |  |  |
| 17 | 0.97 | 1.67 | 0.43 | 1.02 | 0.27 | 0.16 | 0.11 | 0.43 | 0.05 | 0.00 | 0.00 | 0.22 | 0.00 | 0.11 | 0.05 |
| 18 | 2.19 | 3.25 | 1.39 | 1.81 | 0.00 | 0.00 | 0.00 | 0.17 | 0.00 | 0.00 | 0.00 | 0.00 | 0.00 | 0.00 | 0.00 |
| 19 | 0.16 | 1.93 | 5.21 | 5.07 | 0.05 | 0.00 | 0.00 | 1.74 | 0.00 | 0.05 | 0.00 | 0.40 | 0.00 | 0.00 | 0.00 |
| 21 | 0.33 | 0.40 | 0.64 | 0.33 | 0.07 | 0.00 | 0.00 | 0.26 | 0.00 | 0.00 | 0.00 | 0.24 | 0.00 | 0.00 | 0.00 |

N% = ∑ Projection neurons identified within a region / ∑ Total projection neurons identified across both hemispheres × 100

Where cell bodies could be clearly identified, neurons within the injection halo were counted. Furthermore, cell bodies located on boundary lines between areas were only accounted for in the region where the most overlap was present.

**Abbreviations**
17: primary visual cortex
18: second visual cortical area
19: third visual cortical area
20a: temporal visual area a
20b: temporal visual area b
21: fourth visual cortical area
A: A lamina of the lateral geniculate nucleus
A1: A1 lamina of the lateral geniculate nucleus
AMLS: anteromedial lateral suprasylvian visual area
ALLS: anterolateral lateral suprasylvian visual area
C: C lamina of the lateral geniculate nucleus
DLS: dorsal lateral suprasylvian visual cortical area
LP: lateral posterior nucleus of the dorsal thalamus
MIN: medial intralaminar nucleus of the lateral geniculate nucleus
P: perigeniculate lamina of the lateral geniculate nucleus
PMLS: posteromedial lateral suprasylvian visual area
PLLS: posterolateral lateral suprasylvian visual area
PPc: posterior parietal caudal cortical area
PPr: posterior parietal rostral cortical area
PS: posterior suprasylvian visual cortical area
Pul: pulvinar nucleus of the dorsal thalamus
VLS: ventral lateral suprasylvian visual area
VP?: ventral posterior ectosylvian region.

## Results

In the current study BDA tracer injections were made into the occipital visual cortical areas **17**, **18**, **19**, and **21** of the ferret, in order to assess the distribution of both anterograde and retrograde cortico-cortical and cortico-thalamic connections (Figs. 1 – 3). It was evident in all cases that there was more anterograde and retrograde label present in the hemisphere ipsilateral to the injection site than in the contralateral hemisphere, and that this label was present in patches or clusters. Ipsilateral reciprocal connectivity was strongest in the occipital visual areas, with weaker connectivity observed in temporal, suprasylvian and parietal visual areas. Contralateral reciprocal connectivity was observed in the occipital visual areas, with weak connectivity to temporal, suprasylvian and parietal visual areas. Within the cortex ipsilateral to the injection site, the relative occurrence of retrogradely labelled neurons was the greatest within the injected cortical region (Table 1, Fig. 4). Contralaterally, the greatest number of retrogradely labelled neurons was in the homotopic cortical area (Table 1, Fig. 4), except for area **17**, where a stronger connection was found to contralateral area **18**. Ipsilateral reciprocal connectivity was observed throughout the visual nuclei of the dorsal thalamus, and no contralateral connections to the visual thalamus were observed.

**Figure 4:**
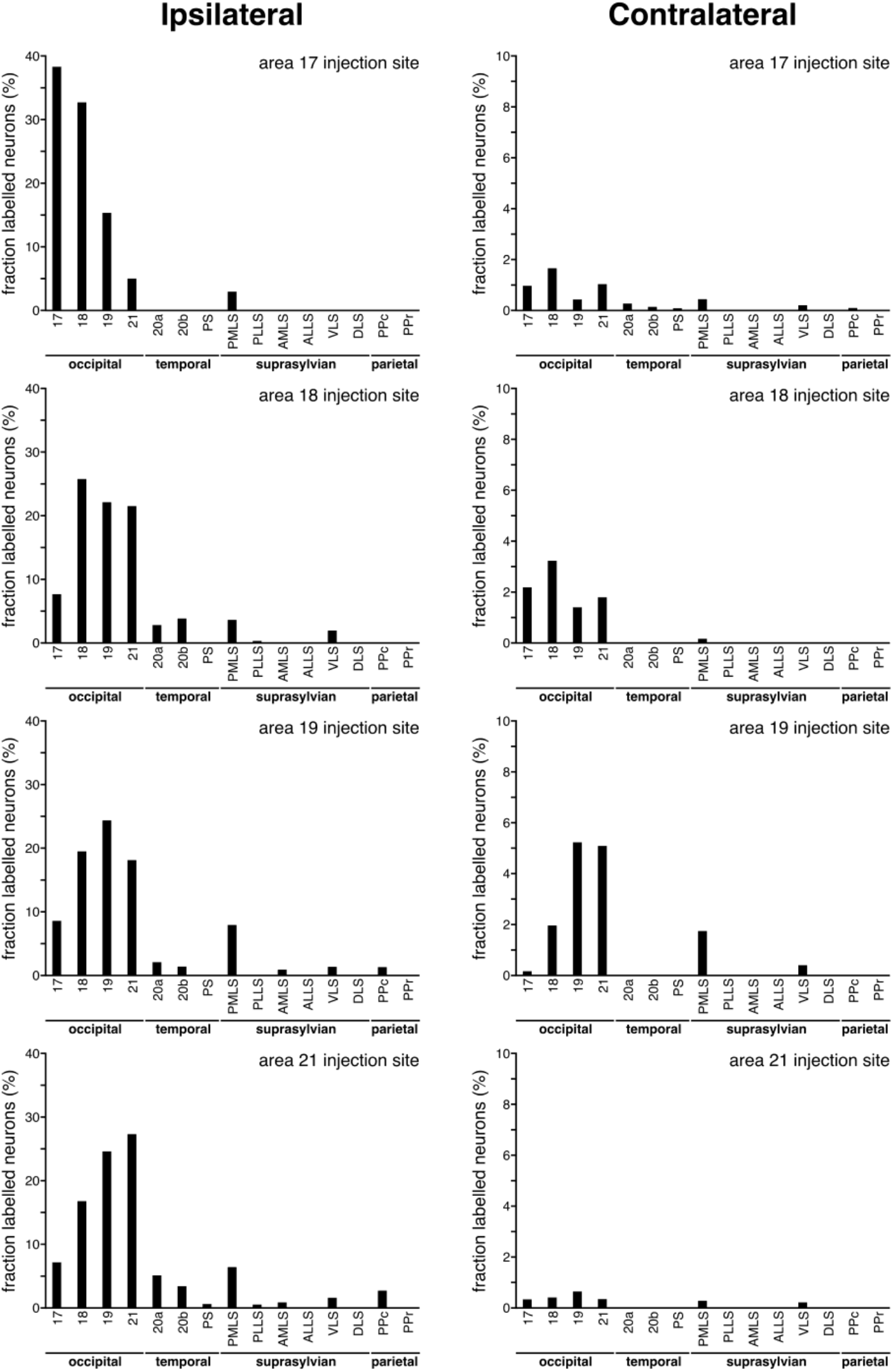
Graphs depicting the quantitative assessment of retrograde connectivity strength within and between cortical areas in the cerebral hemispheres ipsilateral (**left column**) and contralateral (**right column**) to the injection sites made in areas **17** (**top row of graphs**), **18** (**second row of graphs**), **19** (**third row of graphs**) and **21** (**bottom row of graphs**). The values are expressed in percentages, being the fraction of labelled neurons occurring in each cortical area. See list for abbreviations. Note that the majority of retrogradely labelled cells are found within the cortical area injected, and that all occipital areas are connected to each other, both ipsilaterally and contralaterally. Also note than in all cases, both ipsilateral and contralateral, retrogradely labelled cells are found in area **PMLS**, indicating that this cortical area is a node in the visual processing network.

### Connectivity of Area 17

Injection of tracer into area **17** resulted in both ipsilateral anterograde and retrograde label within the occipital visual areas and **PMLS** (Fig. 5). Within area **17** an extensive mediolateral spread of strong anterograde and retrograde labelling was observed (Fig. 5), and this was bordered both medially and laterally by less intense labelling. In area **18** a slightly smaller, but still extensive mediolateral spread of anterograde and retrograde label was observed, with a broader spread of weaker labelling bordering the medial and lateral edges of the more strongly connected region. Strong anterograde and retrograde label was also observed in area **19**, but again the extent of the mediolateral spread was reduced, as was the bordering weaker label which was observed in patches in some places within area **19**. Weak anterograde and retrograde label was observed in the rostral half of area **2**1, where this area shared a border with area **PMLS**. The occasional patch of weak anterograde, or occasional retrogradely labelled cells, was observed in other portions of area **21**. The only cortical area outside of the occipital visual cortex that exhibited connections with area 17 was the suprasylvian area **PMLS**. Within **PMLS**, weak anterograde and retrograde labelling was observed throughout the majority of this cortical area (Fig. 5). These qualitative impressions of the connectivity strength of the different cortical areas with area **17** are supported by the quantification of the relative numbers of retrogradely labelled neurons in the various cortical areas following injection into area **17**. The strongest ipsilateral retrograde connectivity was observed within area **17** (N% = 38.25%), with a decrease in connectivity strength in area **18** (N% = 32.60%), then area **19** (N% = 15.36%), followed by area **21** (N% = 5.01%) and lastly by **PMLS** (N% = 3.13%) (Fig. 4, Table 1).

**Figure 5:**
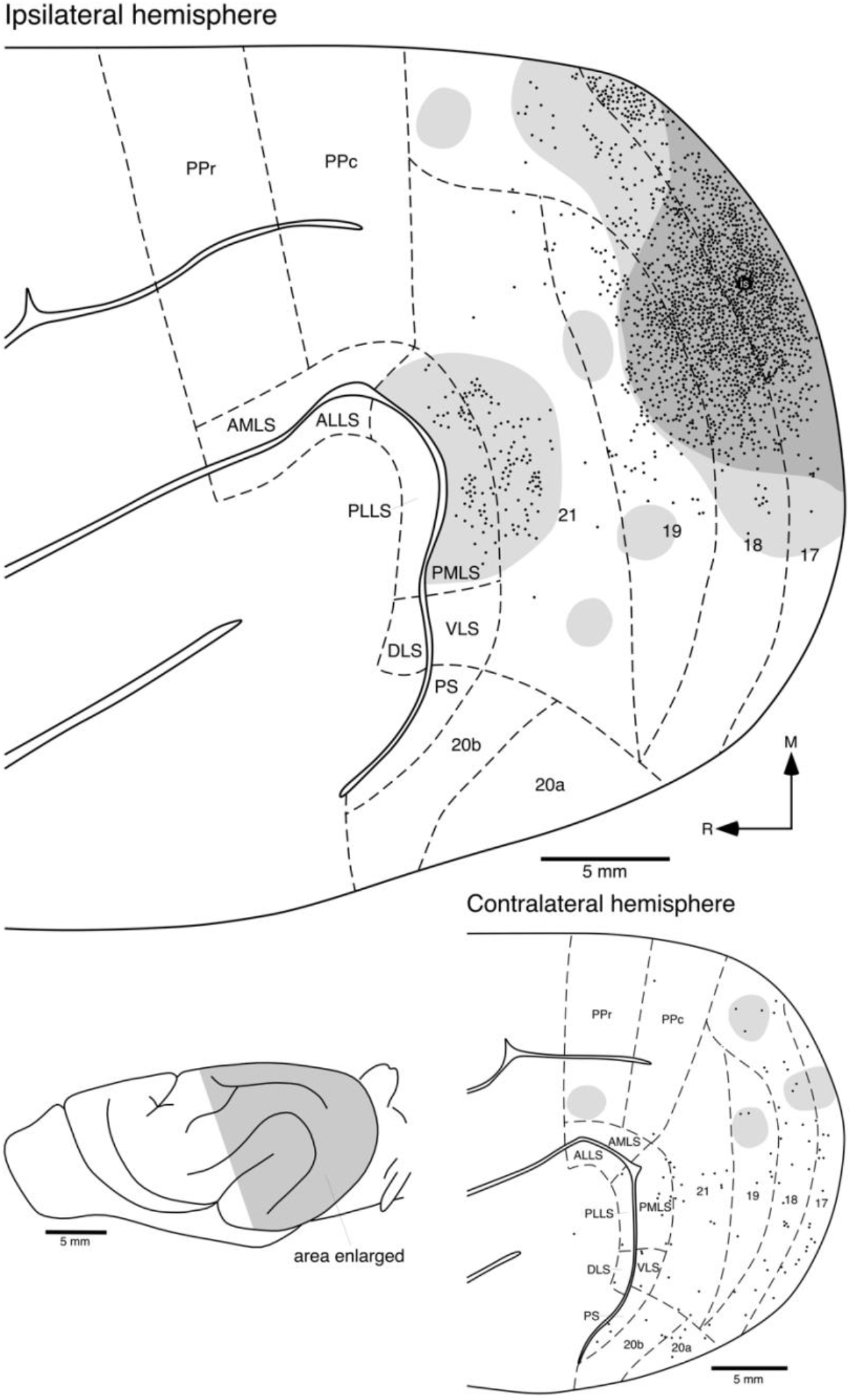
Location of retrogradely labelled cortical neurons (**filled circles**) and anterogradely labelled axons and axon terminals (high density labelling in the **darker grey shading**, low density labelling in the **lighter grey shading**) following transport from the injection site (**is**) located in area **17**. The upper larger image represent the distribution of cells and axons in the ipsilateral semi-flattened caudal half of the cerebral hemisphere, while the lower smaller image represents the semi-flattened caudal half of the cerebral hemisphere contralateral to the injection site. The lateral view of the brain shows the region depicted in the two images (shaded in **light grey**). Note that ipsilaterally, extensive connectivity is seen through the occipital visual areas (**18**, **19** and **21**) and the posteromedial lateral suprasylvian (**PMLS**) visual area. The contralateral connectivity is much weaker, but more widespread. Areal boundaries were demarcated using alternative sections stained for cytochrome oxidase and the boundaries represent approximations based on this stain and available maps of the ferret brain. See list for abbreviations. **M** – medial, **R** – rostral.

Contralateral cortico-cortical connectivity following tracer injection in area 17 was substantially weaker and more diffuse than that observed in the ipsilateral hemisphere (Fig. 5), with a broader spread of retrogradely labelled neurons into cortical areas beyond the occipital areas. In the contralateral area **17** a small patch of weak anterograde label was found homotopic to the injection site, while retrogradely labelled neurons were scattered throughout much of the mediolateral extent of the area. For both areas **18** and **19**, similar connectivity was observed. Within areas **21** and **PMLS**, only diffusely scattered retrogradely labelled neurons were observed. Unlike the ipsilateral connections, in the contralateral hemisphere a small number of retrogradely labelled neurons were observed in the suprasylvian visual areas **PLLS**, **DLS** and **VLS**, and in the temporal visual areas **20a**, **20b** and **PS**. Isolated retrogradely labelled neurons were observed in posterior parietal area **PPc**, while a small patch of anterograde label was observed in **PPr**. Thus, while the connectivity between the hemispheres is far weaker than that within the hemisphere (Figs. 4, 5, Table 1), the contralateral connections are more widespread than the ipsilateral connections.

Following injection of tracer into area **17**, reciprocal connections to the ipsilateral visual nuclei of the dorsal thalamus were observed (Figs. 3a, 6). A small, but dense patch of anterograde and retrograde label within lamina **A/A1** of the lateral geniculate nucleus was seen (Figs. 3a, 6d-i). In addition, small, but dense patches of anterograde projections were observed in the LP nucleus (Figs. 3a, 6b-i) and possibly in the pulvinar nucleus (Fig. 6e). No label was observed in the **C** or **P** lamina of the lateral geniculate nucleus, in the **MIN**, or in the contralateral visual nuclei of the dorsal thalamus.

**Figure 6:**
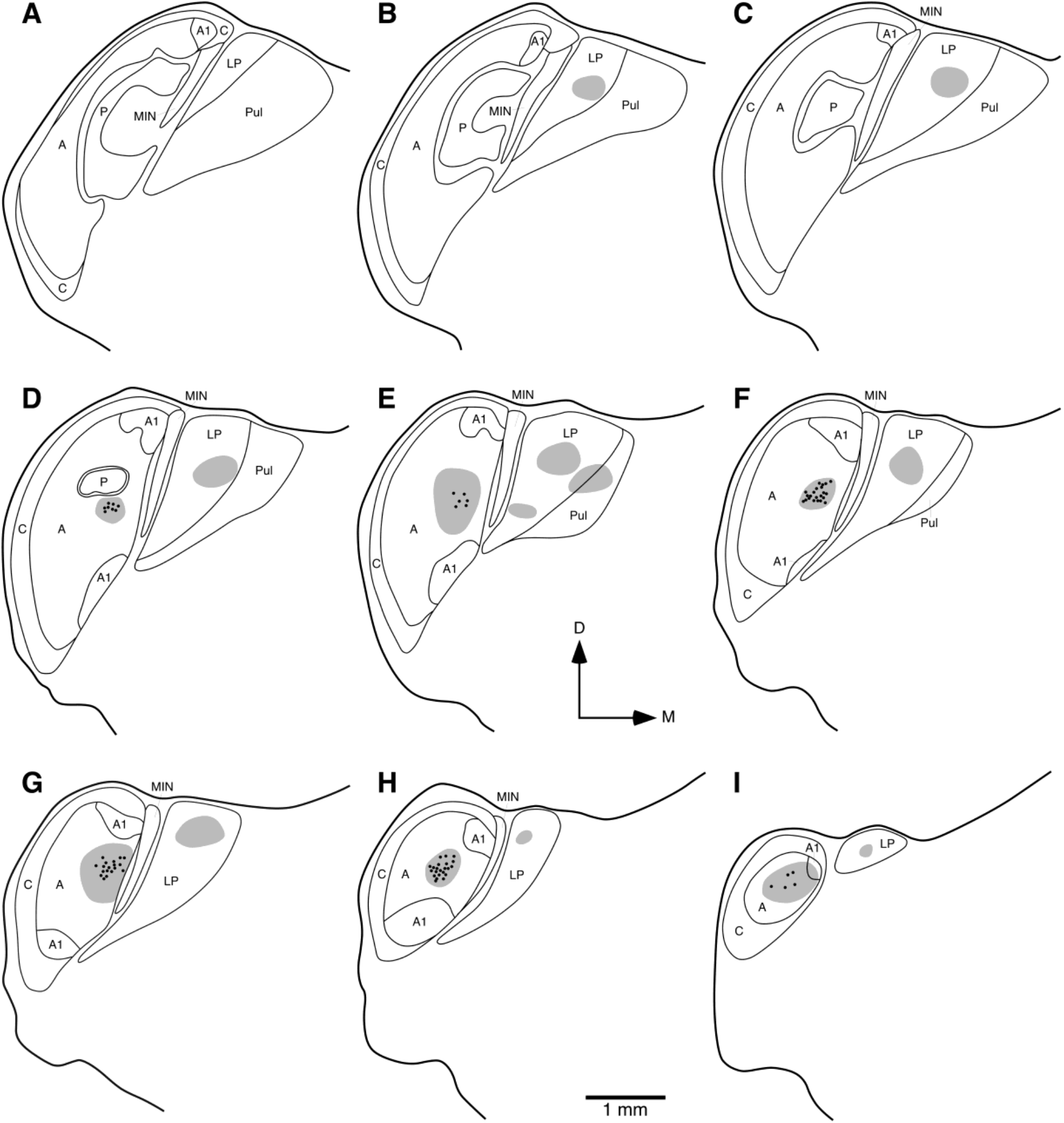
Diagrammatic reconstructions of the location of retrogradely labelled cells (filled circles) and anterogradely labelled axon terminals (high density labelling in the **darker grey shading**, low density labelling in the **lighter grey shading**) in the visual thalamus of the ferret following injection of tracer into the occipital visual area **17**. (**a**) represents the most rostral coronal section, with (**i**) being the most caudal. Each section is approximately 400 μm apart. Note the dense, but restricted, label in the **A** lamina of the lateral geniculate nucleus and the anterogradely labelled region in the **LP** and **Pul**. In all images dorsal (**D**) is to the top and medial (**M**) to the right. See list for abbreviations.

### Connectivity of Area 18

Injection of tracer into area **18** resulted in both ipsilateral anterograde and retrograde label within the occipital, suprasylvian and temporal visual areas (Fig. 7), although the strongest connectivity was observed with other occipital visual areas. As observed following injection of tracer into area **17**, the injections in area **18** resulted in the strongest label being found within this area. Within area **18**, a broad mediolateral spread of anterograde labelling was observed, with the density of this labelling being strongest medial and lateral to the injection site, but with weaker anterograde label being found throughout the majority of the mediolateral extent of the entire area. A similar spread of retrogradely labelled cells was observed in area **18**, although these were not found through the entire area **18**, being somewhat more restricted in their distribution compared to the anterograde label (Fig. 7). The anterograde and retrograde label in area **17**, following injection into area **18**, also showed a dense, but slightly less widespread, band of anterograde and retrograde label in the region immediately caudal to the injection site, with associated mediolateral spread of weaker labelling through much of area **17**. Within area **19**, in the region immediately rostral to the area **18** injection site, a dense band of anterograde and retrograde label was observed, and this was surrounded medially and laterally by weaker labelling that covered the majority of this cortical area. A similar dense band of retrograde labelling, bordered medially and laterally by less dense labelling was observed in area **21**, but the anterograde labelling in area **21** was weaker than that observed in areas **17**, **18** and **19**, less widespread, and did not occupy the entire cortical area (Fig. 7). Within the temporal visual areas, moderately dense retrograde label was observed in areas **20a**, **20b** and **PS**, while weak anterograde label was observed in areas **20a** and **20b**. A few retrogradely labelled neurons were observed in the ventral region of the posterior ectosylvian gyrus (**VP**). In the suprasylvian visual areas **PMLS** and **VLS** a moderate density of retrogradely labelled neurons was observed throughout both cortical areas, and a lower number of labelled cells was observed in area **PLLS**. Weak anterograde label was observed in areas **PMLS**, **PLLS**, **AMLS** and **ALLS**, but not in **VLS** or **DLS** (Fig. 7). The strongest ipsilateral retrograde connectivity was observed within area **18** (N% = 25.94%), with a decrease in connectivity strength in area **19** (N% = 22.18%), then area **21** (N% = 21.58%), followed by areas **17** (N% = 7.80%), 20b (N% = 3.96%), **PMLS** (N% = 3.80%), 20a (N% = 2.99%), with minor connections being observed in areas **PS**, **PLLS**, **VLS** and **AMLS** (Fig. 4, Table 1).

**Figure 7:**
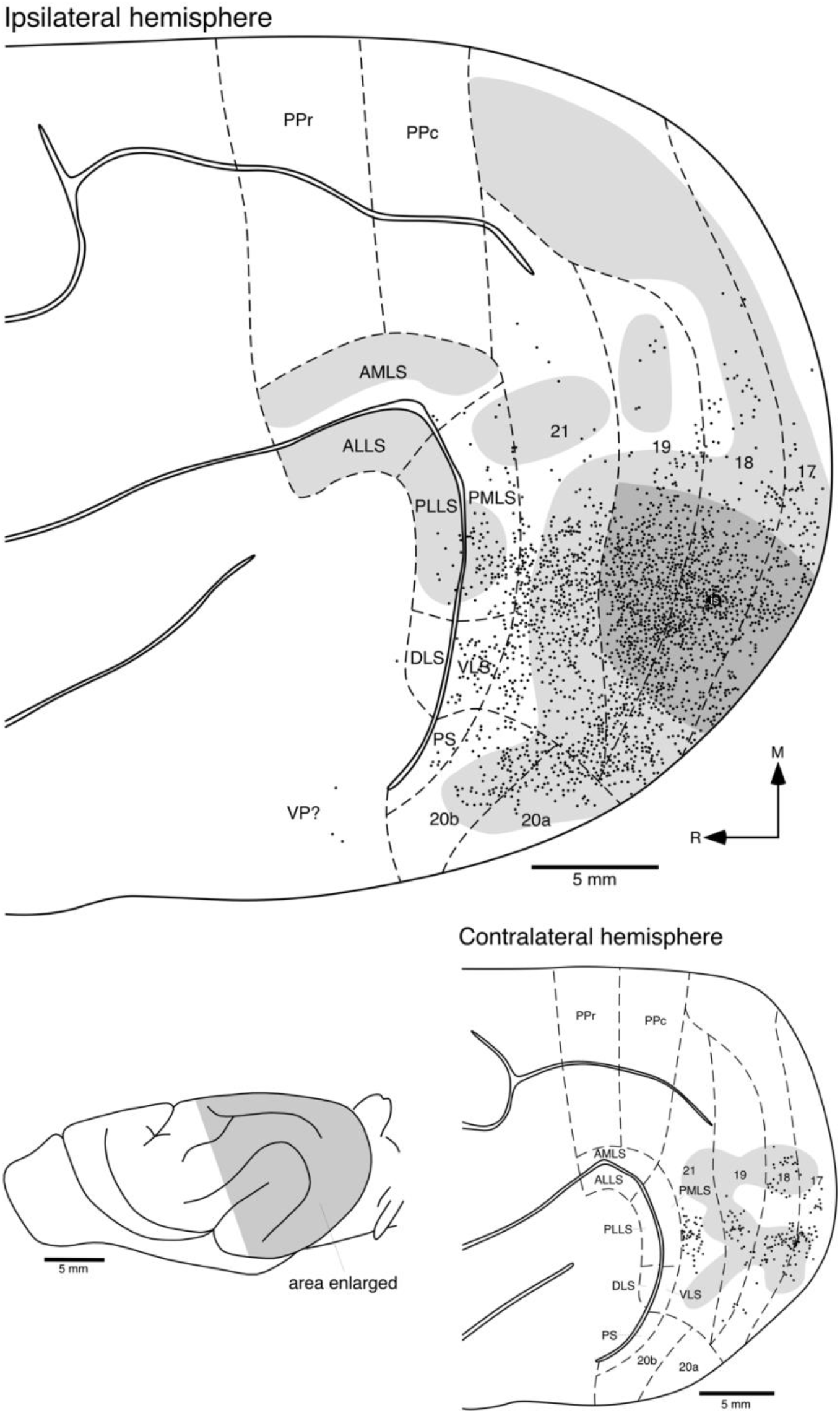
Location of retrogradely labelled cortical neurons (**filled circles**) and anterogradely labelled axons and axon terminals (high density labelling in the **darker grey shading**, low density labelling in the **lighter grey shading**) following transport from the injection site (**is**) located in area **18**. All conventions as provided in the legend to Fig. 5, see list for abbreviations. Note the widespread and extensive ipsilateral connectivity throughout the occipital, temporal and suprasylvian visual areas. The contralateral connectivity, while not as strong as the ipsilateral, is restricted to the occipital visual areas. **VP?** - ventral posterior ectosylvian region.

Contralateral connectivity following injection of tracer into area **18** was restricted to the occipital visual areas **17**, **18**, **19** and **21** (Fig. 7). No contralateral label was observed in the temporal, suprasylvian or parietal visual areas. The highest density of retrogradely labelled cells was observed in contralateral area **18**, in a region homotopic to the injection site, but dorsal to this patch was a second small patch of labelled neurons. Both of these patches of retrogradely labelled cells were contiguous with weak patches of anterograde connectivity. Within the contralateral area **17** the distribution of the retrogradely labelled cells was more contiguous than observed in area **18**, but still closely matched weak patches of anterograde projections. The retrogradely labelled neurons in area **19** were restricted to a patch rostral to the homotopic site of the injection, with the weaker anterograde label being a more mediolaterally spread contiguous region of label. In area **21**, the retrogradely labelled neurons were found in the rostral half of the area in a single moderately dense patch. Interestingly, the contralateral connectivity with area **21** was not reciprocal, as the region where the patch of retrogradely labelled cells was located showed no anterograde projections, which were found to surround this patch medially, laterally and caudally (Fig. 7). Thus, while the connectivity between the hemispheres is far weaker than that within the hemisphere (Figs. 4, 7, Table 1), the contralateral connections are substantially less widespread than the ipsilateral connections following injections of tracer into area **18**.

Following injection of tracer into area **18**, widespread, mostly reciprocal, connectivity was observed throughout all nuclei forming the visual dorsal thalamus (Figs. 3b, 8). Within the lateral geniculate nucleus, retrogradely labelled cells were observed throughout most of the **A/A1** and **C** lamina, as well as the **MIN**, but not in the **P** lamina (Fig. 8). In contrast, dense anterograde labelling, surrounded by weaker labelling, was observed in the **A/AI** and **P** lamina, as well as the **MIN**, but no anterograde labelling was observed in the **C** lamina (Fig. 8). Moderately dense patches of retrogradely labelled cells were observed in both the **LP** and pulvinar nuclei, and these patches were associated with dense anterograde patches of label surrounded by weaker labelling (Figs. 3b, 8a-h). Thus, area **18** is strongly, and mostly reciprocally, connected with all nuclei of the visual portion of the dorsal thalamus of the ferret.

**Figure 8:**
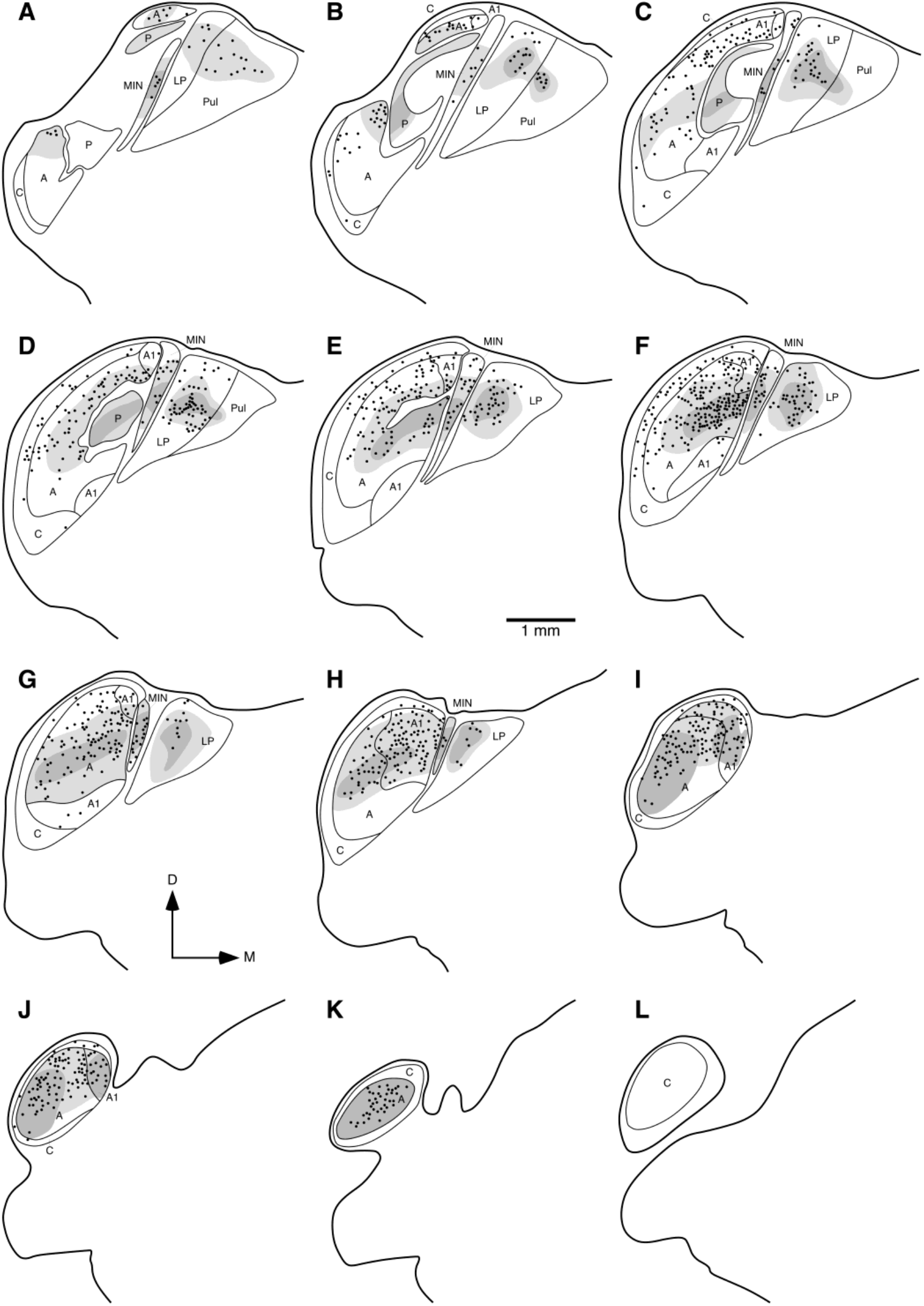
Diagrammatic reconstructions of the location of retrogradely labelled cells and anterogradely labelled axons in the visual thalamus of the ferret following injection of tracer into the occipital visual area **18**.(**a**) represents the most rostral coronal section, with (**l**) being the most caudal. Each section is approximately 400 μm apart. Note the dense, widespread label throughout all subdivisions of the ferret visual thalamus. Conventions as provided in the legend to Fig. 6, see list for abbreviations.

### Connectivity of Area 19

Injection of tracer into area **19** resulted in both ipsilateral anterograde and retrograde label within the occipital, suprasylvian, temporal (including **AEV**), and parietal visual areas, as well as extending into the caudal somatosensory areas and regions of the posterior suprasylvian gyrus (Fig. 9), although the strongest connectivity was observed with other occipital visual areas. The strongest labelling was observed within area **19**, with a high density of retrogradely labelled cells and anterogradely labelled axon terminals found medial and lateral to the injection site, lessening in density with distance from the injection site (apart from a region of strong anterograde label in the lateral most portion of area **19**), but covering the entire cortical area (Fig. 9). In area **18**, retrogradely labelled cells were found throughout much of the mediolateral extent of the cortical area, but were higher in density in a region caudal to the area **19** injection site. The anterograde label in area **18** followed a similar pattern, with a high density of label in the middle of the cortical area, surrounded medially and laterally by weaker label. In addition, the lateral most part of area **18** had a high density patch of anterograde label (Fig. 9). Similarly, in area **17** there was a region of high density of retrogradely labelled cells, with lower labelled cell densities medial and lateral to this region. Interestingly, unlike in areas **19** and **18**, the region of high density of anterograde label in area **17** was not matched to the region of high retrograde label, which was found medial to the region of high labelled cell density. Again, this region of high anterograde label was found bordered medially and laterally by regions of less dense label, but this covered the majority of area **17**. Rostral to the area **19** injection site, retrogradely labelled cells were observed throughout the mediolateral extent of area **21**, with a higher density being observed closer to the injection site. A small patch of dense anterograde label was observed in area **21** near to the area **19** injection site, but the remainder of area **21** was in receipt of a lower density anterograde projection. Retrogradely labelled neurons were found in the temporal visual areas **20a**, **20b** and **PS**, but the numbers of cells labelled in **PS** was lower than that observed in **20a** and **20b**. Interestingly, area **20a** was not in receipt of any anterograde projections following the area **19** injection, but a dense patch of anterograde connectivity was observed crossing the border between area **20b** and **PS**, with light patches of anterograde label being observed in area **AEV** and the posterior ectosylvian gyrus (Fig. 9). Within the suprasylvian visual areas, retrogradely labelled neurons were observed in areas **AMLS**, **PMLS** and **VLS**, with the greatest density of these being observed in **PMLS**. In addition, less dense anterograde label was observed in all suprasylvian areas except **DLS**, and dense anterograde label was observed in **PMLS** in the same regions of this area where a high density of retrogradely labelled cells were observed. A low density of retrogradely labelled neurons were observed in the lateral most aspect of **PPc**, while in the lateral most aspects of **PPc** and **PPr** light anterograde label was also observed. In addition, patches of anterograde label were observed in the medial half of **PPc** and **PPr** (Fig. 9). Lastly, weak anterograde labelling was observed in the caudal somatosensory areas **SII** and **SIII**. The strongest ipsilateral retrograde connectivity was observed within area **19** (N% = 24.32%), with a decrease in connectivity strength in area **18** (N% = 19.38%), then area **21** (N% = 18.06%), followed by areas **17** (N% = 8.56%), **PMLS** (N% = 7.92%), with minor connection strengths being observed in areas **20a**, **20b**, **PS**, **PPc**, **AMLS** and **VLS** (Fig. 4, Table 1).

**Figure 9:**
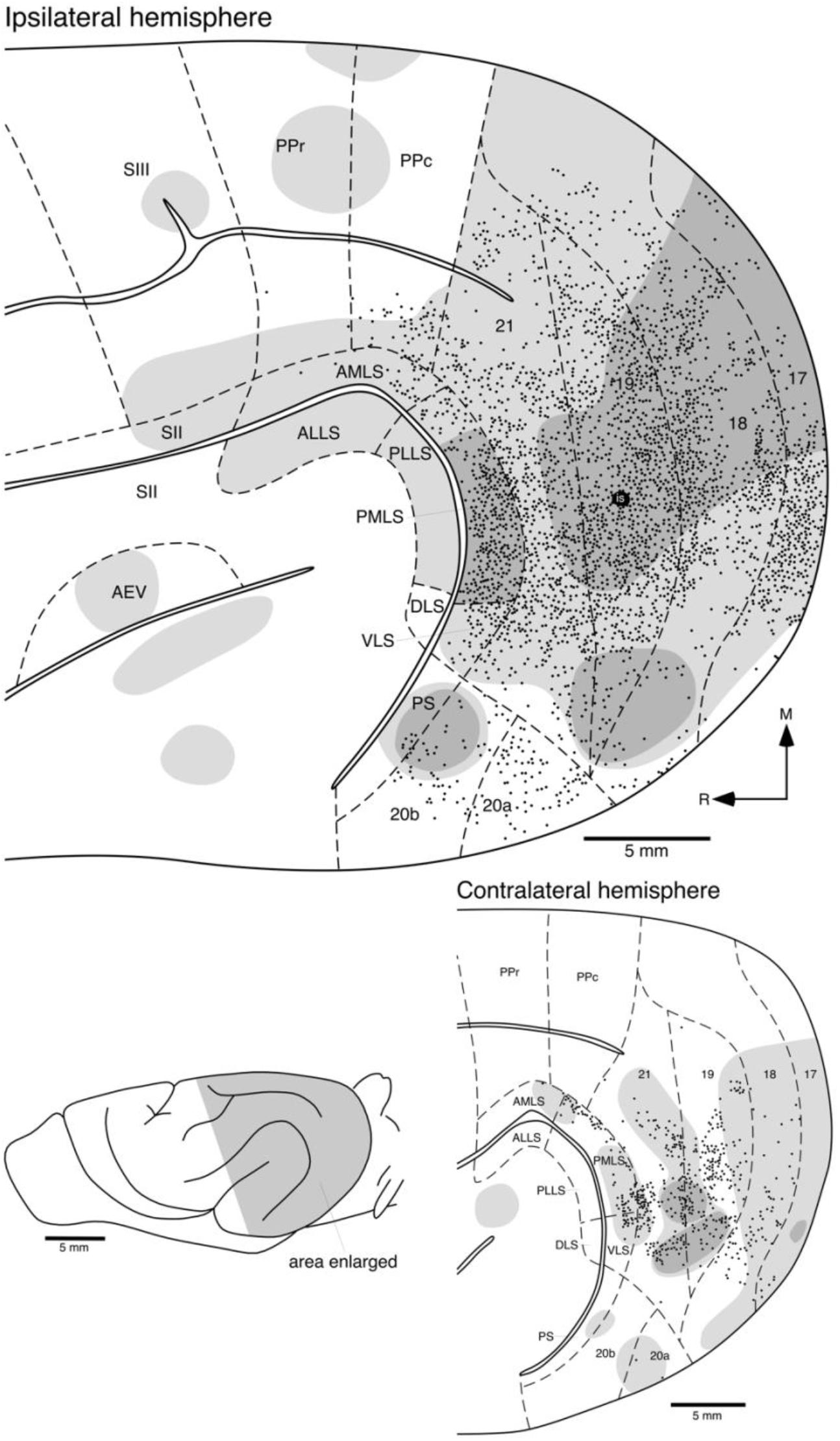
Location of retrogradely labelled cortical neurons (**filled circles**) and anterogradely labelled axons and axon terminals (high density labelling in the **darker grey shading**, low density labelling in the **lighter grey shading)** following transport from the injection site (**is**) located in area **19**. All conventions as provided in the legend to Fig. 5, see list for abbreviations. Note the widespread ipsilateral connectivity throughout the occipital, temporal (including **AEV**, the anterior ectosylvian visual area), suprasylvian and parietal visual areas. Anterograde connectivity was also observed in the second (**SII**) and third (**SIII**) somatosensory areas, as well as higher order areas ventral to the **AEV** and auditory cortex. Contralateral connections are observed in the occipital, temporal (including **AEV**) and suprasylvian visual areas.

Contralateral connectivity following injection of tracer into area **19** was found in the occipital, temporal and suprasylvian visual areas (Fig. 9), with no contralateral label observed in the parietal visual areas or somatosensory areas. The strongest patch of retrogradely labelled cells was observed in contralateral area **19**, in a region homotopic to the injection site, but spreading throughout the entire area. Similarly, a dense patch of anterograde label was found in area **19** in a region homotopic to the injection site, and spreading across much of the contralateral area **19**, although the anterograde and retrograde connections are not entirely overlapping (Fig. 9). Within the contralateral area **18** a low density of retrogradely labelled cells was observed throughout the lateral half of this area and for the most part was contiguous with low density anterograde label. Very few retrogradely labelled cells were observed in the contralateral area **17** following injection into area **19**, but a low density anterograde projection was observed throughout much of this cortical area. Low density patches of anterograde projections were observed in areas **20a**, **20b** and **PS**, with a patch near the medial border of area **AEV**, but only a very few callosally projecting labelled cells were observed in area **20a** (Fig. 9). Within the suprasylvian visual areas, moderate densities of retrogradely labelled cells were observed in areas **PMLS** and **VLS**, with a few cells in **AMLS**. These cell densities were matched with low density anterograde projections in **PMLS**, **VLS** and **AMLS** (Fig. 9). Again the connectivity between the hemispheres is far weaker than that within the hemisphere (Figs. 4, 9, Table 1), and the contralateral connections are substantially less widespread than the ipsilateral connections following injections of tracer into area **19**.

Following injection of tracer into area **19**, widespread, mostly reciprocal, connectivity was observed throughout all nuclei forming the visual dorsal thalamus (Figs. 3c, 10), although the strength of the reciprocal connection to the pulvinar was weak. Within the lateral geniculate nucleus, retrogradely labelled cells were observed in the **A/A1**, **C** and **P** lamina, as well as the **MIN** (Fig. 10). Similarly, dense anterograde labelling, surrounded by weaker labelling, was observed in the **A/AI**, **C** and **P** lamina, as well as the **MIN** (Fig. 10). Moderately dense patches of retrogradely labelled cells were observed in the **LP** nucleus, and these patches were associated with dense anterograde patches of label surrounded by weaker labelling (Figs. 3c, 10a-g). Only a minor reciprocal connection was observed between area **19** and pulvinar nucleus. Thus, area **19** is strongly, and mostly reciprocally, connected with most of the nuclei of the visual portion of the dorsal thalamus of the ferret.

**Figure 10:**
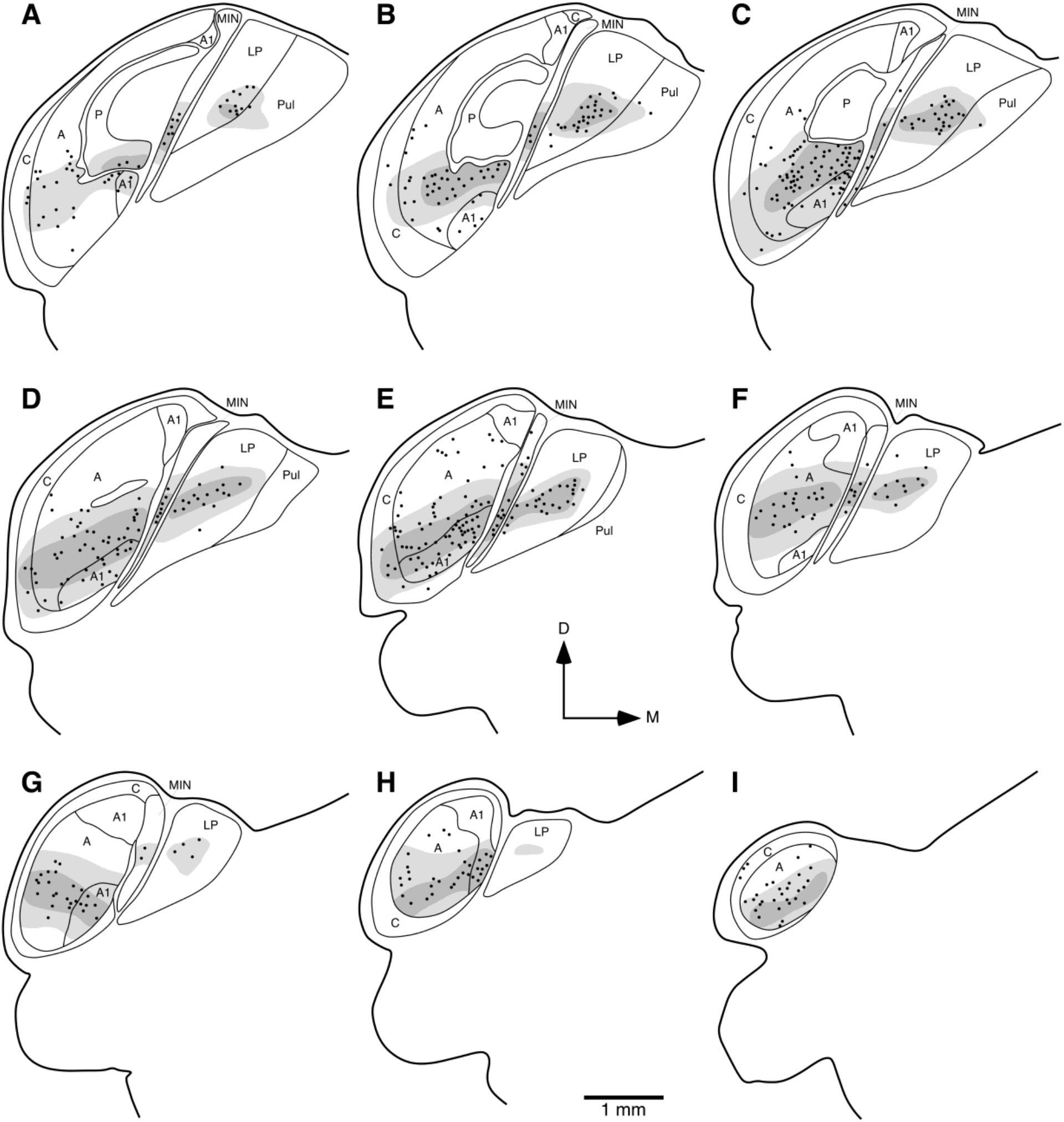
Diagrammatic reconstructions of the location of retrogradely labelled cells and anterogradely labelled axons in the visual thalamus of the ferret following injection of tracer into the occipital visual area **19**. (**a**) represents the most rostral coronal section, with (**i**) the most caudal. Each section is approximately 400 μm apart. Note the dense, widespread label throughout all subdivisions of the ferret visual thalamus apart from the pulvinar nucleus. Conventions as provided in the legend to Fig. 6, see list for abbreviations.

### Connectivity of Area 21

Injections into area **21** led to substantial and widespread ipsilateral connectivity throughout all occipital visual regions, all temporal areas including the **AEV** and the ventral portion of the posterior ectosylvian gyrus, suprasylvian and posterior parietal visual areas, although the strongest connectivity was with other occipital visual areas (Fig. 11). As with injections in all the other occipital visual areas, the injection in area **21** showed the strongest connectivity within the same area. A high to moderate density of retrogradely labelled cells were found throughout area **21**, and this was matched by high densities of anterograde projections, although these were weaker at the medial and lateral borders of the area. Area **19** showed a similar pattern of connectivity, with a high to moderate density of retrogradely labelled cells found throughout area **19**, being matched mostly by a high density of anterograde projections (Fig. 11). The density of retrogradely labelled cells in area **18** was slightly lower than that observed in areas **21** and **19**, and were not found throughout area **18** with the medial and lateral edges of area **18** lacking retrogradely labelled cells. A smaller patch of high density anterograde tracer was observed in area **18**, caudal to the area **21** injection site, and the density of the anterograde label decreased in both a medial and lateral direction. A similar pattern was observed in area **17**, but the mediolateral extent of the higher density anterograde projection was greater in area **17** than area **18** (Fig. 11). Retrogradely labelled cells were found throughout the temporal visual areas, being moderately dense in areas **20a**, **20b** and **PS**, and less dense in area **AEV** and the posterior ectosylvian gyrus. These regions were, for the most part, matched by a low density anterograde projection, which was more extensive in area **AEV** than the labelled cells, and was found in the ventrocaudal portion of the posterior ectosylvian gyrus ventral to the suprasylvian visual area **DLS** (Fig. 11). All suprasylvian visual areas evinced retrogradely labelled neurons, but these were found in the highest densities in areas **PMLS** and **VLS**. Areas **PMLS** and **VLS** were in receipt of a high density of anterograde projections, with **AMLS**, **ALLS**, **PLLS** and **DLS** receiving low density anterograde projections (Fig. 11). A moderate density of retrogradely labelled neurons were observed in the most lateral aspect of area **PPc**, with a few cells being observed in the medial portion of PPc and the lateral portion of **PPr**. Both the medial and lateral portions of **PPc** were in receipt of low density patches of anterograde projections, while these were limited to the lateral aspect of **PPr**. The strongest ipsilateral retrograde connectivity was observed within area **21** (N% = 27.25%), with a decrease in connectivity strength in area **19** (N% = 24.45%), then area **18** (N% = 16.77%), followed by areas **17** (N% = 7.13%), **PMLS** (N% = 6.31%), with minor connection strengths being observed in other areas as described above (Fig. 4, Table 1).

**Figure 11:**
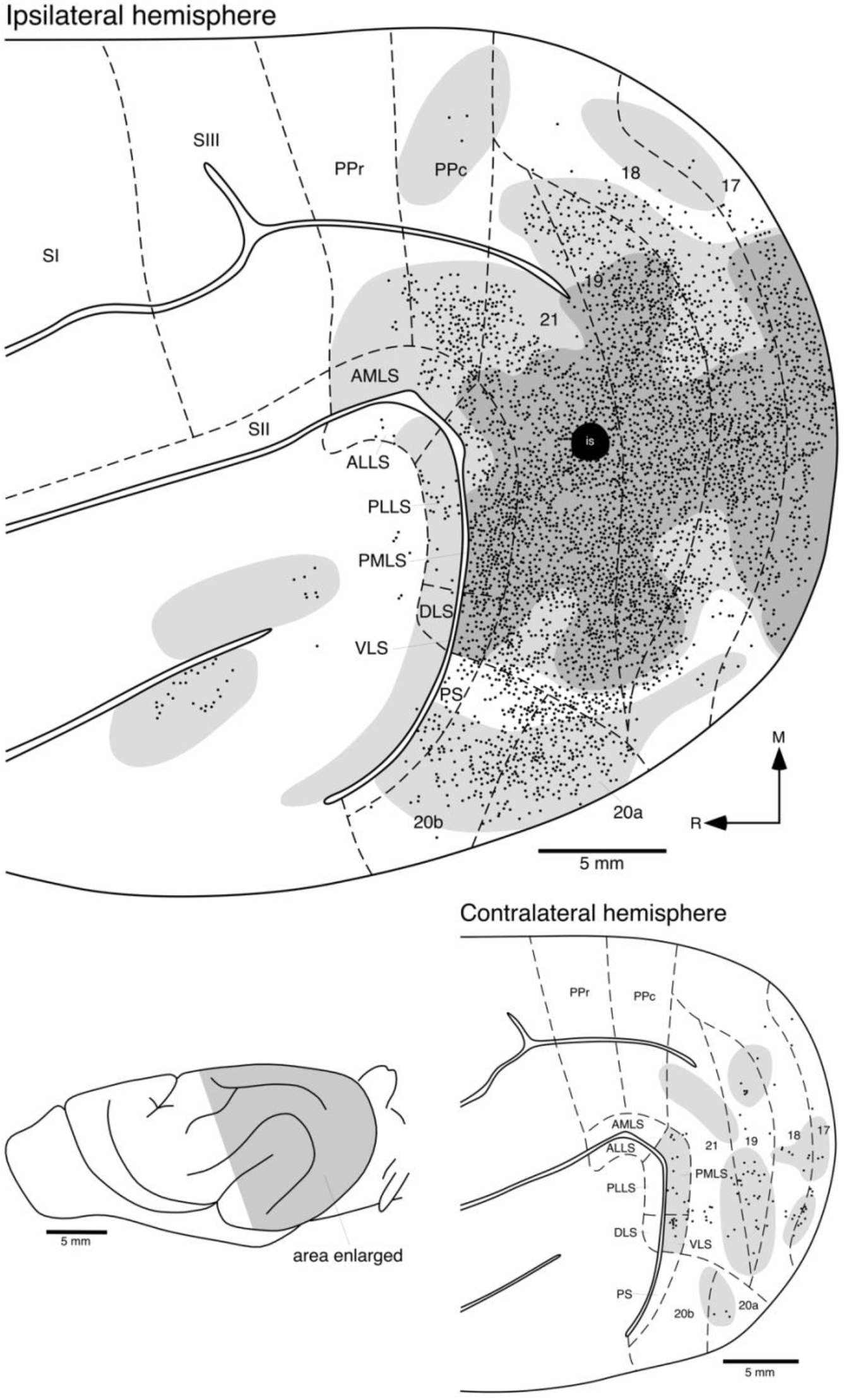
Location of retrogradely labelled cortical neurons (**filled circles**) and anterogradely labelled axons and axon terminals (high density labelling in the **darker grey shading**, low density labelling in the **lighter grey shading**) following transport from the injection site (**is**) located in area **21**. All conventions as provided in the legend to Fig. 5, see list for abbreviations. Note the widespread ipsilateral connectivity throughout the occipital, temporal (including **AEV**, the anterior ectosylvian visual area), suprasylvian and parietal visual areas. Anterograde connectivity was also observed in the higher order areas ventral to the **AEV** and auditory cortex. Contralateral connections are observed in the occipital, temporal and posterior suprasylvian visual areas. **SI** – primary somatosensory cortex.

Contralateral connectivity following injection of tracer into area **21** was found in the occipital, temporal and suprasylvian visual areas **PMLS** and **VLS** only (Fig. 11). The strongest patch of retrogradely labelled cells was observed in contralateral area **19**, not in the homotopic region of the injection site in area **21**, however, retrogradely labelled cells were found in area **21** in a low density. Patches of low density anterograde label were found in both areas **21** and **19** in regions matching the location of the retrogradely labelled cells and somewhat beyond (Fig. 11). Within the contralateral areas **18** and **17** low densities of retrogradely labelled cells were observed in the lateral half of these areas and were contiguous with low density anterograde label. A few retrogradely labelled neurons were observed in area **20a**, but not in other temporal visual areas, and these were found in the same region as a light patch of anterograde label that extended a small way into area **20b**. A higher number, although low in density, of retrogradely labelled cells were observed in areas **PMLS** and **VLS** and both these areas were in receipt of a low density anterograde projection. Again the interhemispheric connectivity is far weaker than that within the hemisphere (Figs. 4, 11, Table 1), and the contralateral connections are substantially less widespread than the ipsilateral connections following injections of tracer into area **21**.

Following injection of tracer into area **21**, widespread, mostly reciprocal, connectivity was observed throughout all nuclei forming the visual dorsal thalamus, except for the pulvinar nucleus which showed no connectivity with area **21** (Figs. 3d, 12). Within the lateral geniculate nucleus, retrogradely labelled cells were observed in the **A/A1** and **C** lamina, as well as the **MIN** (Fig. 12), but not the **P** lamina. Similarly, dense anterograde labelling, surrounded by weaker labelling, was observed in the **A/AI** and **C** lamina, as well as the **MIN** (Fig. 12). A moderately dense patch of retrogradely labelled cells were observed in the **LP** nucleus, and this patch was associated with a dense anterograde patch of label surrounded by weaker labelling (Figs. 3d, 12a-j). Thus, area **21** is strongly, and mostly reciprocally, connected with many of the nuclei of the visual portion of the dorsal thalamus of the ferret.

**Figure 12:**
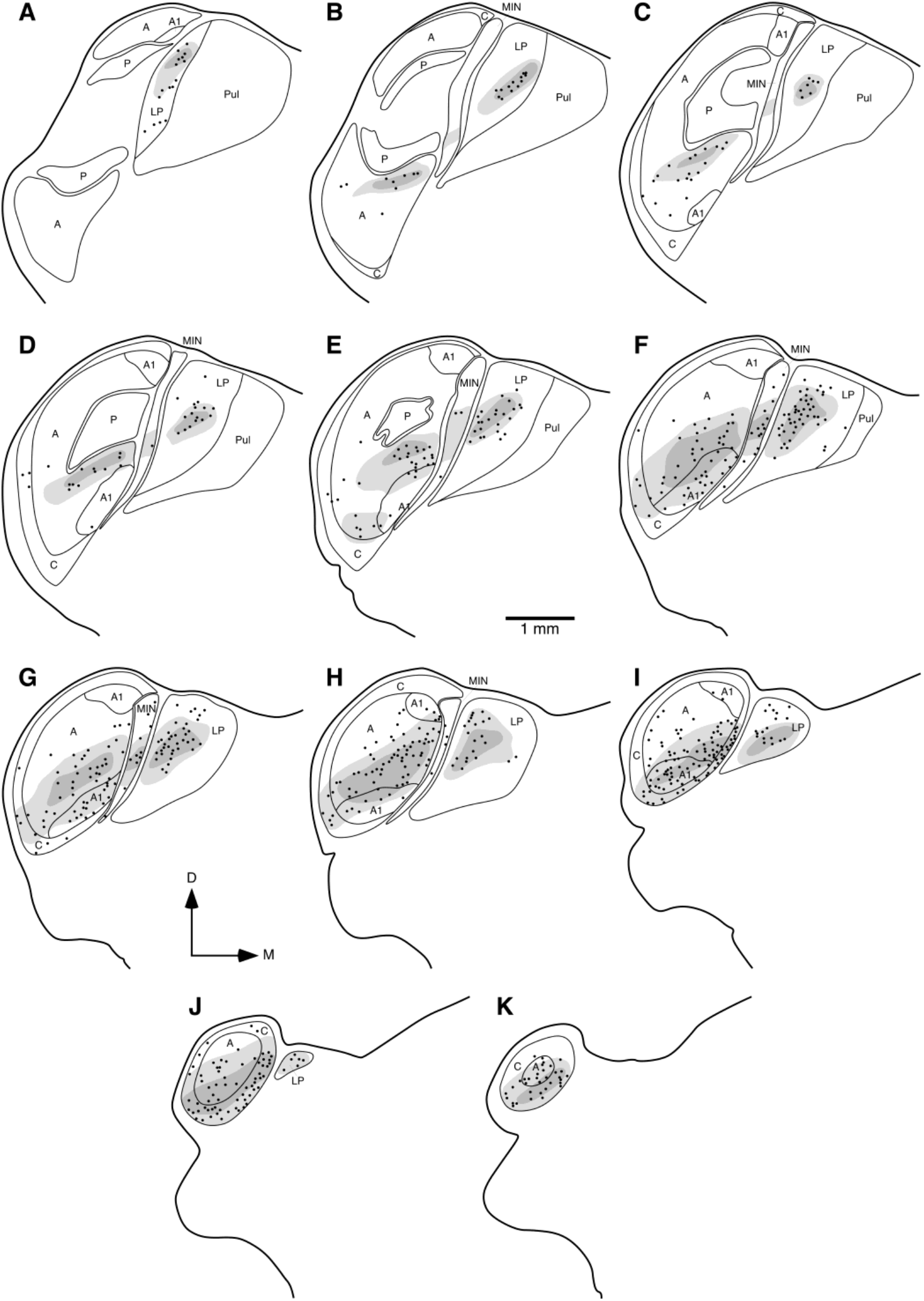
Diagrammatic reconstructions of the location of retrogradely labelled cells and anterogradely labelled axons in the visual thalamus of the ferret following injection of tracer into the occipital visual area **21**.(**a**) represents the most rostral coronal section, with (**k**) being the most caudal. Each section is approximately 400 μm apart. Note the dense, widespread label throughout all subdivisions of the ferret visual thalamus apart from the pulvinar nucleus. Conventions as provided in the legend to Fig. 6, see list for abbreviations.

## Discussion

The present study details the ipsilateral and contralateral cortico-cortical and cortico-thalamic connectivity of the occipital visual areas **17**,**18**, **19** and **21** in the ferret using standard anatomical tract-tracing methods. These data will contribute to the Ferretome (www.ferretome.org) collation of ferret macroscopic brain connectivity, to studies of cortical connectivity in relation to other aspects of brain structure and function as well as studies of cortical connectivity across mammals.

### General observations

Although areas **17**, **18**, **19** and **21** are strongly interconnected ipsilaterally, the number of ipsilateral areas to which a specific area was connected increased, as a gradient, from 4 areas connected with area **17**, 12 with area **18**, 13 with area **19**, and 15 with area **21** (Fig. 13). This finding corroborates previous observations in cortical connectivity studies in other mammalian species (Beul & Hilgetag, 2015). The connectivity with the cortical areas of the contralateral hemisphere exhibited a similar pattern, but the connectivity strength was substantially lower and more restricted in terms of the breadth of connected areas. This observation is in agreement with connectional studies of the macaque prefrontal cortex as well as mouse and rat cortices (Barbas, Hilgetag, Saha, Dermon, & Suski, 2005; Goulas, Uylings, & Hilgetag, 2017; Swanson, Hahn, & Sporns, 2017), whereby the contralateral hemisphere maintains a subset of the cortico-cortical projections observed in the ipsilateral hemisphere (Swanson et al., 2017). Numerous connectional similarities and differences were observed when comparing the occipital visual areas of the ferret and the cat.

**Figure 13:**
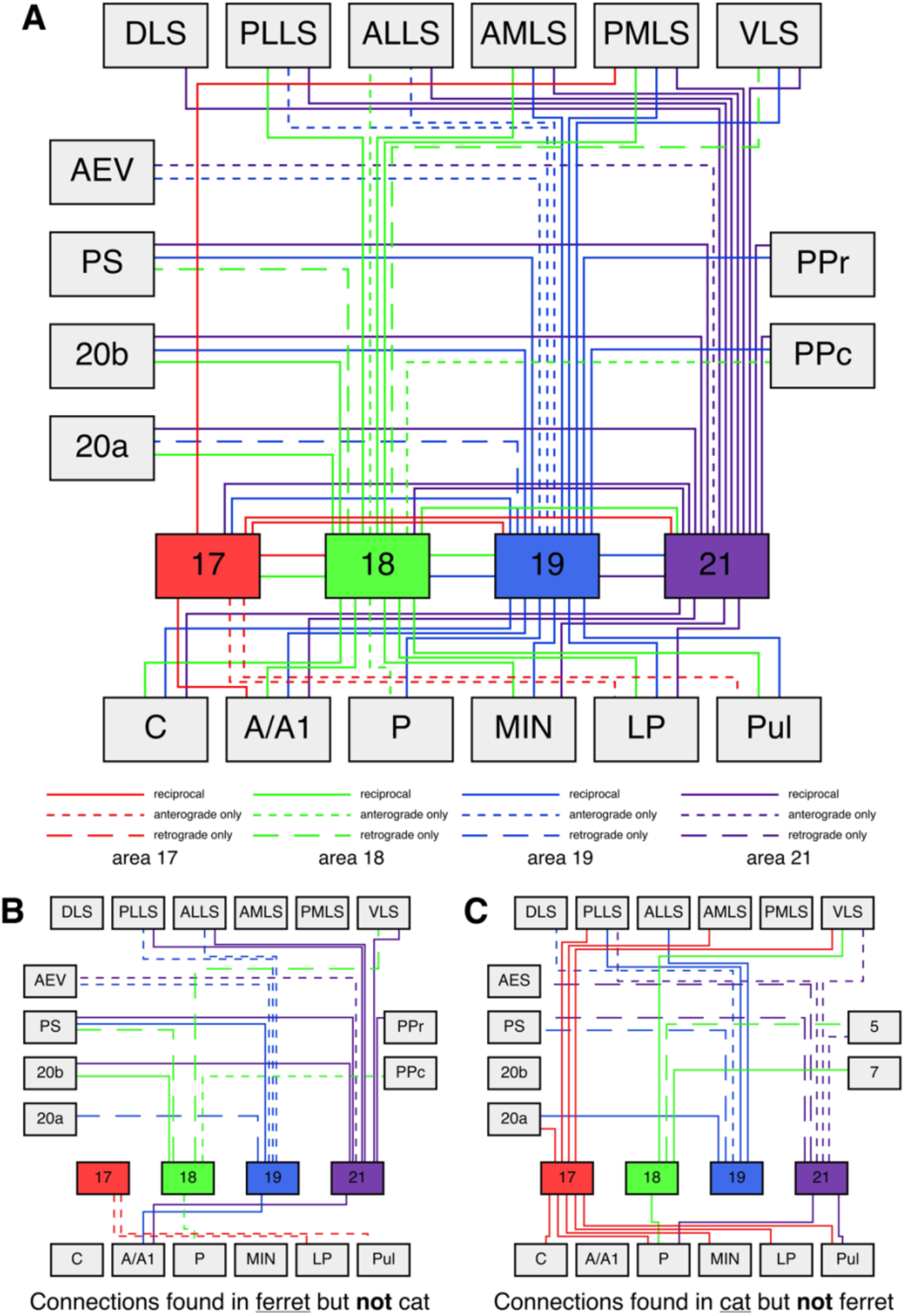
Wiring diagrams depicting the connectivity of areas **17**, **18**, **19** and **21** with each other, other visual cortical areas and the visual thalamus in the ferret (**a**), the connections of these areas observed in the ferret but not the cat (**b**), and the connections of these areas overs in the cat but not the ferret (**c**). Each colour represents a specific cortical area (**17** – red, **18** – green, **19** – blue, **21** – purple), with solid lines representing reciprocal connections, short dashed lines anterograde connections only, and long dashed lines retrograde connections only. (**a**) Of the potential 252 connections (for each cortical area 21 potential reciprocal, anterograde only or retrograde only connections) of these four areas to the different visual cortical and thalamic regions, a total of 70 connections are found in the ferret, indicating the high connectivity of the visual system in this species. Areas **19** and **21** appear to show the broadest connectivity, while area **PMLS** appears to be a connection hub located outside the occipital cortex, with reciprocal connections to all occipital cortical areas. (**b**) In the ferret 21 connections not found in the cat were observed, while (**c**) in the cat 25 connections not found in the ferret were observed, many of which involve area **17** (Heath & Jones 1970, 1971; Symonds & Rosenquist, 1984; Scannell et al., 1995).

### Area 17 connectivity – ferret vs cat

Area **17** in the ferret displays limited inter-areal ipsilateral connectivity, being reciprocally connected with areas **18**, **19**, **21** and **PMLS**. While area **17** of the cat exhibits these same reciprocal connections, additional reciprocal connections with areas **PLLS**, **AMLS**, **VLS** and **20a** have been observed in the cat (Heath & Jones, 1970, 1971; Symonds & Rosenquist, 1984; Scannell et al., 1995) (Fig. 13). Although much weaker than the ipsilateral connectivity, the contralateral, or callosal connectivity of the ferret reached a broader range of areas. Reciprocal callosal connectivity was observed for areas **17**, **18** and **19**, while retrograde connectivity was observed in areas **21**, **20a**, **20b**, **PS**, **PMLS**, **PLLS**, **DLS**, **VLS** and **PPc**, and anterograde connectivity was observed in area **PPr**. Similarly, in the cat, reciprocal callosal connectivity was observed for areas **17**, **18**, and **19**, with additional connections being observed in areas **AMLS** and **PMLS**. In contrast to the ferret, area **21** displayed callosal anterograde connectivity with area **17** in the cat (Sanides, 1978; Segraves & Rosenquist, 1982a,b; Segraves & Innocenti, 1985). Concerning the connectivity of area **17** with the visual nuclei of the dorsal thalamus, area **17** in the ferret was observed to be reciprocally connected with the **A** lamina of the **LGN**, and sends projections (anterograde connectivity) to the **LP** and pulvinar. In contrast, area **17** in the cat is reciprocally connected with all laminae of the **LGN** (**C**, **A**, **P** and **MIN**) and the **LP** and pulvinar (Symonds, Rosenquist, Edwards, & Palmer, 1981; Scannell, Burns, Hilgetag, O’Neil, & Young, 1999) (Fig. 13). Thus, when comparing the connectivity of area **17** between the ferret and the cat, it is clear that area **17** of the cat is more broadly connected than that of the ferret.

### Area 18 connectivity – ferret vs cat

Area **18** of the ferret exhibits a much broader ipsilateral inter-area connectivity pattern than area **17**, being reciprocally connected with areas **17**, **19**, **21**, **20a**, **20b**, **PMLS**, **PLLS** and **AMLS**, sending anterograde connections to areas **ALLS** and **PPc**, and receiving projections from areas **VLS** and **PS** (Fig. 13). While cat area **18** shares many of these connections, unlike the ferret it receives input from area **5** (**PPr** of the ferret), is reciprocally connected with areas **VLS** and **7** (**PPc** of the ferret), and lacks connectivity with areas **20b** and **PS** observed for the ferret (Heath & Jones, 1970, 1971; Symonds & Rosenquist, 1984; Scannell et al., 1995). Area **18** of the ferret exhibited more restricted contralateral connectivity compared to the ipsilateral connectivity, evincing reciprocal callosal connections only with areas **17**, **18**, **19** and **21**. In contrast, the cat displayed reciprocal callosal connections with the occipital visual areas as well as **AMLS** and **PMLS**. In addition, callosal retrograde connections were identified with **20a**, while anterograde callosal connectivity was observed in area **VLS** (Sanides, 1978; Segraves & Rosenquist, 1982a,b; Segraves & Innocenti, 1985). Area **18** of the ferret presents with reciprocal connections to all portions of the visual thalamic nuclei (all laminae of the **LGN** and the **LP** and pulvinar nuclei), apart from the **P** lamina of the **LGN**, where area **18** projects to this lamina. In contrast, area **18** of the cat exhibits reciprocal connectivity with all portions of the visual thalamic nuclei, including the **P** lamina of the **LGN** (Symonds et al., 1981; Scannell et al., 1999). Thus, while there are some minor differences in the connectivity patterns between area **18** of the ferret and cat, the broad patterns are very similar and it cannot be stated that one species has a more broadly connected area **18** than the other.

### Area 19 connectivity – ferret vs cat

Area **19** of the ferret exhibited a slightly broader ipsilateral inter-area connectivity pattern than area **18** (Fig. 13), and was reciprocally connected with area **17**, **18**, **21**, **PMLS**, **AMLS**, **20b**, **PS**, **PPc** and **PPr**, projected to areas **PLLS**, **ALLS** and **AEV**, and received projections from area **20a**. Area **19** of the cat shares the vast majority of these connections, but distinct to ferret area **19**, area **19** of the cat projects to area **DLS**, but does not project to area **AES** (**AEV** of the ferret). In addition, the connections of area **19** in the cat with areas **PLLS**, **ALLS** and **20a** are reciprocal, whereas they are only unilateral in the ferret, while area **19** of the cat receives projections from area **PS**, yet ferret area **19** is reciprocally connected with **PS** (Heath & Jones 1970, 1971; Symonds & Rosenquist, 1984; Scannell et al., 1995) (Fig. 13). Area **19** of the ferret exhibited far broader contralateral connectivity than area 18, being reciprocally connected with areas **17**, **18**, **19**, **21**, **PMLS**, **AMLS**, **VLS** and **20a**, and projecting to areas **20b** and **PS**. Similarly, area **19** in the cat presented with the most extensive callosal connectivity, with connectivity extending beyond the areas callosally connected in the ferret. Reciprocal callosal connectivity was observed in areas **17**, **18**, **19**, **21**, **AMLS** and **PMLS**, while projections to areas **20a**, **20b**, **ALLS**, **PLLS**, **DLS** and **VLS** were identified (Heath & Jones, 1970; Segraves & Rosenquist, 1982a,b). Area **19** of the ferret revealed reciprocal connections to all portions of the visual thalamic nuclei (all laminae of the **LGN** and the **LP** and pulvinar nuclei), with all of these connections being observed in the cat, apart from the reciprocal connection to lamina **A** of the **LGN**, which is absent in the cat (Symonds et al., 1981; Scannell et al., 1999) (Fig 13). Thus, as with area **18**, area **19** of the ferret and cat are connected in a very similar manner, with minor variations mostly associated with anterograde or retrograde connectivity, but the distinct lack of a reciprocal connection of cat area **19** with the **A** lamina of the **LGN** (the largest and most cell dense retino-recipient lamina of the **LGN**), which is observed in the ferret, indicates the potential for functional differences between areas **19** of the ferret and cat which may be elucidated with single cell recordings.

### Area 21 connectivity – ferret vs cat

Of the four occipital visual areas studied, area **21** evinced the broadest ipsilateral inter-areal connectivity. We observed that area **21** of the ferret was reciprocally connected with areas **17**, **18**, **19**, all six suprasylvian visual areas, **20a**, **20b**, **PS**, **PPc** and **PP**r, while also projecting to area A**EV**. Area **21** of the cat is also connected with all these areas, but the connections with areas **PLLS**, **VLS** and **5** (**PPr** of the ferret) are anterograde only, while those with **PS** and **AES** (**AEV** of the ferret) are retrograde only (Heath & Jones 1970, 1971; Symonds & Rosenquist, 1984; Scannell et al., 1995) (Fig. 13). Area **21** of the ferret exhibited a similar, although more restricted, pattern of callosal connectivity as area **19**, being reciprocally connected with areas **17**, **18**, **19**, **21**, **PM**LS, **VLS**, **20a** and **20b**. Area **21** of the cat has a similar pattern of callosal connectivity as that seen in the ferret. Reciprocal callosal connections were identified in areas **18**, **19**, **21**, **PMLS** and **20b**. Within the contralateral hemisphere, cat area **21** also projected to areas **20a**, **DLS** and **PLLS**, while receiving projections from areas **17** and **VLS** (Segraves & Rosenquist, 1982a,b). Area **21** of the ferret was observed to have reciprocal connections with lamina **C**, **A** and **MIN** of the **LGN**, as well the **LP** nucleus, but lacked connections with lamina **P** of the **LGN** and the pulvinar nucleus. In contrast, while cat area **21** shares most of these connections, it lacks connections with the **A** lamina of the **LGN** seen in the ferret, but is reciprocally connected with the **P** lamina of the **LGN** and the pulvinar nucleus (Symonds et al., 1981; Scannell, et al., 1999), both connections not observed in the ferret (Fig. 13). While the pattern of connectivity of area **21** in the ferret and the cat is very similar, and the variances are mostly attributable to whether a connection is anterograde or retrograde, it is striking that the ferret area **21** does not receive input from the pulvinar nucleus, and that the cat area **21** does not receive input from lamina **A** of the **LGN**, again indicating potential functional differences in visual processing by these areas between species.

### Similarities in connectivity patterns of occipital visual areas of the ferret and the cat

While differences between connectivity patterns between the species are of interest, before these are discussed, it must be emphasized that the inter-area ipsilateral and contralateral cortico-cortical and corticothalamic connections observed in the current study in the ferret and in previous studies in the cat (Heath & Jones 1970, 1971; Symonds et al., 1981; Symonds & Rosenquist, 1984; Scannell et al., 1995, 1999; Symonds et al., 1995), are essentially very similar. The connectivity patterns outlined in Figure 13 indicate that each cortical area injected has the possibility to project to 21 different sites (15 ipsilateral cortical areas, 6 thalamic sites, callosal connectivity not included in this figure), meaning that in the four occipital visual areas in the ferret and cat, when looking at site to site projections (eliminating such additional variables as cortical layers and potential subdivisions of the **LP** and pulvinar nuclei), up to 84 site to site connections could exist in each species. Our current study of the ferret identified a total of 66 out of the 84 possible connections (78.6%), indicating a high degree of interconnectivity within the visual system of the ferret. For the cat, 70 out of the 84 possible projections (83.3%) have been identified in previous studies (Heath & Jones, 1970, 1971; Symonds et al., 1981; Symonds & Rosenquist, 1984; Scannell et al., 1995, 1999), which, given biological variability, indicates that the general degree of connectivity of the occipital visual areas in the ferret and cat is highly similar. Interestingly, of the 66 connections found in the ferret, 7 were not found in the cat, while of the 70 connections reported for the cat, 11 are not found in the ferret. Additionally, 12 of the connections differed in type between the species (being either reciprocal, anterograde or retrograde in one species, and different in the other species). Thus, for the four occipital visual areas in the ferret and cat, 47 connections (out of a theoretical possibility of 84) to other cortical areas or the visual dorsal thalamus are the same, with 12 connections varying in type but not connected sites, and 18 being species specific (7 for the ferret, 11 for the cat). This high degree of connectional similarity matches the similarity in the areal organization of the visual cortical areas between ferret and cat (Symonds et al., 1981; Symonds & Rosenquist, 1984; Scannell et al., 1995, 1999; Manger et al., 2002a,b, 2004), potentially indicating an order specific baseline for the organization of the cortical areas, visual thalamic nuclei and connectivity amongst these within the Carnivora (Manger, 2005). Further studies on other carnivore species will be needed to confirm this potential order specificity.

The present study, and similar previous studies in the cat (Heath & Jones, 1970, 1971; Symonds et al., 1981; Symonds & Rosenquist 1984; Scannell et al., 1995, 1999) indicate that the four occipital visual areas are strongly interconnected, and likely form a similar network for the flow of information through the occipital region. In addition, the present study and those in the cat indicate that external to the occipital visual areas, area **PMLS** forms a distinct network hub, as it is reciprocally connected with all four occipital visual areas, distinguishing **PMLS** from all other non-occipital visual cortical regions. **PMLS** is involved in detection of visual motion perception, as well as motion analysis, shifts in attention and the discrimination of speed (Spear & Baumann, 1975; Rauschecker, von Grünau, & Poulin, 1987; Krüger, Kiefer, Groh, Dinse, & von Seelen, 1993; Cantone et al., 2006; Rokszin, Márkus, Braunitzer, Berényi, Benedek, & Nagy, 2010) and thus it is heavily implicated in dorsal visual stream functioning, which involves the parietal cortex (Cantone et al., 2006; Cloutman, 2013). Furthermore, the notion exists that **PMLS** may have a role in molding neuronal responses of the visual cortex, as well as facilitating the presence of parallel circuits to carry out different visual functions (Spear & Baumann 1979; Cantone et al., 2006). The concept that **PMLS** is a specific network hub in the ferret and cat visual system is supported by the evidence that not all occipital visual areas are directly connected to the posterior parietal cortex and when these connections do exist, they are not strong (Table 1). However, **PMLS** and the posterior parietal cortex are directly connected (Cantone et al., 2006), implicating **PMLS** as the gateway, or network hub, to the parietal cortex for the dorsal stream (Cloutman, 2013).

### Differences in connectivity patterns of occipital visual areas of the ferret and the cat

While the similarities in cortical areas, visual thalamic nuclei and patterns of connectivity are very similar in the ferret and cat, certain differences of interest do emerge. Of these differences, it is the variation in the degree of connectivity to the visual nuclei of the dorsal thalamus, especially the laminae of the **LGN** that appear to be the most salient. In the ferret, area **17** is reciprocally connected with only lamina **A** of the **LGN** and projects to the **LP** and pulvinar nucleus, whereas in the cat area **17** is reciprocally connected with all four components of the **LGN** as well as the **LP** and pulvinar nuclei (Symonds et al., 1981; Scannell et al., 1999). Areas **18** and **19** are reciprocally connected to all 6 divisions of the visual thalamic nuclei in both ferret and cat, while area **21** in the ferret is reciprocally connected to lamina **A**, **C** and **MIN** of the **LGN** and the **LP** nucleus, whereas in cat it is connected to all six subdivisions of the visual dorsal thalamus (Symonds et al., 1981; Scannell et al., 1999). Thus, we can conclude that the information flow from the dorsal thalamus, and likely in the thalamocortical loop, is more heavily influenced by the thalamus in the cat compared to the ferret. In addition, it appears that this distinction is strongest for area **17** of the cat (Fig. 13). In contrast, it appears that for the ipsilateral inter area cortical connectivity patterns, area **21** of the ferret is more strongly interconnected than area **21** of the cat (Fig. 13). This difference in connectivity patterns appears to place more emphasis on neuronal analysis of information earlier in the visual cortical network in the cat (**LGN** and area **17**), whereas the emphasis appears to be later in the visual cortical network in the ferret (area **21**).

Such a difference in processing emphasis or strength may be reflective of anatomical, behavioural, developmental or phylogenetic differences between the two species. Cats have brains approximately 6 times larger than those of ferrets (30 g vs 5 g, Bininda-Emonds, Gittleman, & Purvis, 1999). In addition, the cat has frontalized eyes allowing for an increased frontal binocular field compared the ferret (Morgan, Henderson, & Thompson, 1987; Herrera & Mason, 2007; Larsson, 2015). Lastly, the last common ancestor of the ferret and cat occurred over 50 million years ago and belong to different suborders within the Carnivora (Bininda-Emonds et al., 1999; Agnarsson, Kuntner, & May-Collado, 2010). Expansive binocular vision allows cats to be stalk, ambush and pounce predators. Thus, the emphasis of connectivity on the primary portions of the visual cortical network may allow the cat to more effectively use binocular summation, lowering the detection threshold for a stimulus from potential fast moving prey, allowing more effective extraction of salient stimuli from the environment (Cartmill, 1992; Heesy, 2009; Martin, 2009; Larsson, 2015). In contrast, ferrets dig and burrow, scavenge, and eat small vertebrates, carrion, insects, and occasionally fruit and honey (Evans & An, 1998; Horner & Biknevicius, 2010). Thus, a great emphasis on the extraction of more complex visual features of the environment would appear to be more salient to the ferret, this being reflected in the connectivity emphasis shift towards higher order cortical areas.

The differential in connectivity patterns between the two species may be reflective of different connectivity developmental profiles, where early in development exuberant connectivity is similar between species, but processes associated with the development of connections, such as synaptic pruning, may lead to the observed differences between species (Innocenti, 1995, 2017). Alternatively, given the greater than 50 million years since the most recent common ancestor of the ferrets and cats, the differences observed may be phylogenetic in nature rather than adaptive to the animal’s life history. Future studies on other carnivore species from the two major carnivore lineages, of similar detail to those undertaken on the ferret and cat, are needed in order to understand whether the observed differences are based in species-specific adaptive mechanisms, or are reflective of their independent phylogenetic histories.

## Acknowledgements

The authors wish to thank Mrs. Sonata Valentiniene for her consistently high-quality histological preparations.

## Role of Authors

PRM and GI designed the study and undertook the experimental aspects of the study. LAD, CCH and PRM analyzed the material and LAD wrote the first draft of the paper, which was subsequently edited by GMI, CCH and PRM. All authors had full access to all of the data in the study and take responsibility for the integrity of the data and the accuracy of the data analysis.

